# Beyond the reach of homology: successive computational filters find yeast pheromone genes

**DOI:** 10.1101/2021.09.28.462209

**Authors:** Sriram Srikant, Rachelle Gaudet, Andrew W. Murray

## Abstract

The mating of fungi depends on pheromones that mediate communication between two mating types. Most species use short peptides as pheromones, which are either unmodified (e.g., α-factor in *Saccharomyces cerevisiae*) or C-terminally farnesylated (e.g., **a**-factor in *S. cerevisiae*). Peptide pheromones have been found by genetics or biochemistry in small number of fungi, but their short sequences and modest conservation make it impossible to detect homologous sequences in most species. To overcome this problem, we used a four-step computational pipeline to identify candidate **a**-factor genes in sequenced genomes of the Saccharomycotina, the fungal clade that contains most of the yeasts: we require that candidate genes have a C-terminal prenylation motif, are fewer than 100 amino acids long, contain a proteolytic processing motif upstream of the potential mature pheromone sequence, and that closely related species contain highly conserved homologs of the potential mature pheromone sequence. Additional manual curation exploits the observation that many species carry more than one **a**-factor gene, encoding identical or nearly identical pheromones. From 332 fungal genomes, we identified strong candidate pheromone genes in **238** genomes, covering **13** clades that are separated from each other by at least 100 million years, the time required for evolution to remove detectable sequence homology. For one small clade, the *Yarrowia*, we demonstrated that our algorithm found the **a**-factor genes: deleting all four related genes in the **a**-mating type of *Yarrowia lipolytica* prevents mating.

## Introduction

Fungi are an ancient lineage of eukaryotes whose extant members shared a common ancestor more than one billion years ago [1]. Within a species, the mating depends on signaling by diffusible pheromones between haploid cells of two different mating types (Figure 1A) [2]. One important group is the *Saccharomycotina*, which emerged roughly 400 million years ago and contains most of the yeasts: unicellular fungi that lack fruiting bodies. Many yeast species play key roles as model systems, in the food industry, and as crop and human pathogens. Finding the pheromones of economically important yeast will help to understand and control their mating as well as shed light on the evolution of these molecules and the receptors that recognize them.

**Figure 1.**
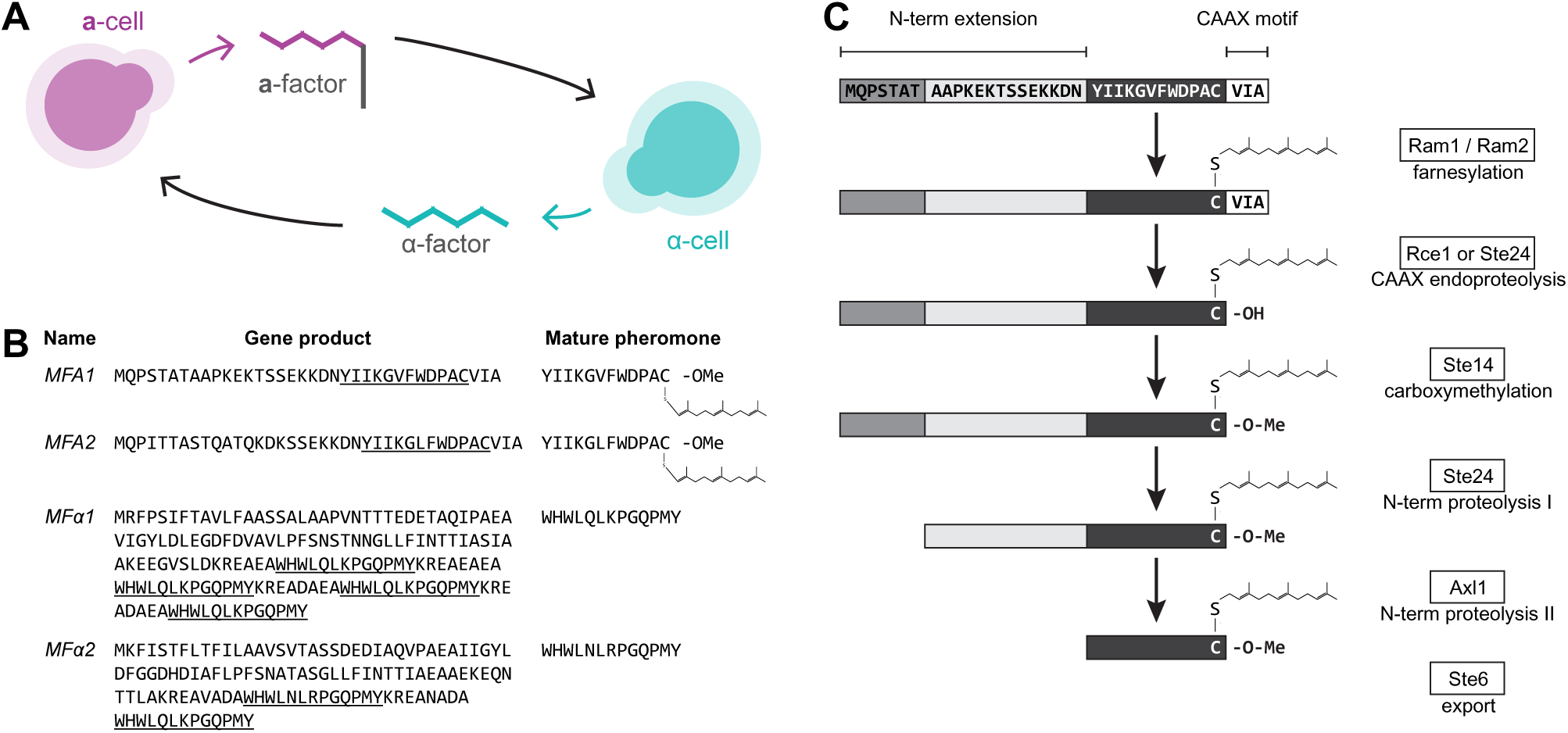
*S. cerevisiae* pheromones are produced by cleaving and modifying precursor peptides. (A) Haploid *S. cerevisiae* have two mating types, **a** and α. Their mating with each other is initiated by the secretion of diffusible peptide pheromones that are recognized by a G protein-coupled receptors (GPCR): **a**-cells (magenta) secrete the lipidated peptide pheromone **a**-factor, which is recognized by the **a-**factor receptor expressed on α-cells, while α-cells (cyan) secrete the peptide pheromone α-factor, which is recognized by the α-factor receptor expressed by **a**-cells. (B) Mating pheromones (underlined) are encoded within small precursor peptides by the *MFA1* and *MFA2* genes (for **a**-factor) and *MFα1* and *MFα2* genes (for α-factor). These peptides require several maturation steps before their secretion as biologically active molecules. (C) The modifications of initial products of the *MFA1* and *MFA2* genes that produce **a**-factor. Broadly, there are two stages of maturation, C-terminal modifications (S-thiol farnesylation, -AAX proteolysis (white bars), and carboxymethylation), followed by two steps of N-terminal proteolysis (two grey bars). The mature pheromone (black bar) is then exported from the cytosol through a dedicated ABC transporter, Ste6.

The best studied mating system is that of the baker’s yeast *Saccharomyces cerevisiae*. Genetic screens and biochemistry have identified pheromones, the proteins that produce and export them, pheromone receptors, and their associated signaling pathways. The two pheromones in *S. cerevisiae* are the 13-amino acid peptide α-factor produced by α-cells and **a**-factor, the 12-amino acid C-terminally farnesylated peptide produced by **a**-cells (Figure 1B). Multiple copies of α-factor are present in each of two mating factor genes (*MFα1* and *MFα2*) and are released by proteolysis in the endoplasmic reticulum allowing the pheromone to be exported in secretory vesicles [3, 4]. In contrast, **a**-factor is encoded in two loci (*MFA1* and *MFA2*) as a precursor peptide that undergoes maturation in the cytosol to produce a single pheromone molecule (Figure 1C) [5, 6]. **a**-factor maturation involves C-terminal farnesylation and carboxymethylation, followed by N-terminal proteolysis. These steps occur in the cytoplasm, after which **a**-factor is pumped across the plasma membrane by Ste6, a member of the ABC family of ATP-dependent transporters. In *S. cerevisiae*, maturation is essential for bioactivity; unmodified pheromone is trapped in the cytosol because it is not recognized and exported by Ste6 [5, 7].

The Saccharomycotina fungi lie within the Ascomycota clade and the distinction between an unmodified α-factor and a farnesylated **a**-factor is maintained in the Ascomycota. In the Basidomycota, the sister clade to the Ascomycota, all mating pheromones are farnesylated, suggesting that the ancestral mating pheromones were farnesylated, poorly soluble molecules [8]. The peptide sequences of both **a**-like and α-like pheromones diverge across fungal species with cognate receptors and transporters co-evolving to maintain activity [9–11].

The first fungal pheromones were identified as lipidated peptides by biochemical isolation from the extracellular medium of basmidomycetes *Rhodosporidium toruloides* [12, 13] and *Tremella mesenterica* [14, 15]. Since then, pheromones have been biochemically isolated and characterized in laboratory yeasts, filamentous fungi and crop pathogens by isolation from extracellular medium [2, 8, 16, 17] but this approach depends on discovering the environmental conditions that stimulate mating and obtaining sufficient material to determine the pheromone’s peptide sequence.

An alternative route to pheromone identification is to look for homologs of known pheromones in sequenced fungal genomes. Because pheromones are small peptides and are only modestly conserved this approach only works over a limited phylogenetic distance. In principle, this problem could be solved by exploiting conservation in the order of genes along a chromosome (synteny): looking for orthologs of the longer genes on either side of the *S. cerevisiae* **a**-factor and then searching, more sensitively, for a pheromone ortholog that lies in the stretch of DNA between these two genes. Unfortunately, synteny is less conserved than pheromone sequence. A search for homologs of one of the *S. cerevisiae* genes that encode **a**-factor, *MFA2*, finds hits in two clades that diverged roughly 100 million years ago from the clade that enjoyed a whole genome duplication and includes *S. cerevisiae*: in the nearer clade (which contains *Zygosaccharomyces rouxii* and *Torulospora delbrueckii*) the **a**-factor homologs retain their syntenic location, but in the further one (which contains *Kluyvermoyces lactis, Lachancea waltii* and *Eremothecium gossypii*) synteny has been lost [18].

We built a computational pipeline to identify farnesylated fungal pheromones in sequenced genomes. Our algorithm successively discards sequences that lack the C-terminal signal sequence for farnesylation, lack a sequence that controls proteolytic processing, and lack strongly conserved homologs in closely related species. Manual curation privileges sequences that occur more than once in the same genome and identifies all the known pheromones in the Saccharomycotina. Overall, we find known or candidate pheromone genes in 238 of 332 mined genomes. To test the ability to identify novel pheromones, we verified that deletion of all copies of the candidate farnesylated pheromone of *Yarrowia lipolytica* (whose last common ancestor with *S. cerevisiae* existed more than 300 million years ago) prevented mating of the **a**-cells of this species.

## Results

### Algorithmic sieve to find candidate pheromone open reading frames (ORFs)

Given the limitations of syntenic methods and the constraints on homology search for small unstructured peptides, we designed a new approach to identify farnesylated peptide pheromones from sequenced genomes using successive identification of conserved features of previously identified pheromones across the fungal lineage as a series of sieves to identify candidate pheromone sequences.

Biogenesis of **a**-factor in *S. cerevisiae* involves a number of essential post-translation modifications by dedicated enzymes [5] (Figure 1C). We assume that these conserved enzymes play a role in all fungal pheromone maturation, and therefore our algorithm identifies candidate pheromone genes by searching for the sequence motifs that these enzymes recognize. The post-translational maturation can be broadly separated into two parts – C-terminal farnesylation and carboxy-methylation, followed by N-terminal proteolytic cleavage (Figure 1C).

In eukaryotes, prenylation is achieved by a conserved enzyme complex recognizing a conserved sequence motif [8]. The enzyme complex (Ram1/Ram2) prenylates by S-thiol addition of a farnesyl group (a C15 isoprenoid) to the cysteine in the C-terminal sequence CAAX, with C representing the modified cysteine, A being small aliphatic residues and X being a small amino acid.

In addition to the S-farnesyl modification, the pheromones need to be carboxymethylated for bioactivity [19, 20]. The C-terminal AAX residues of *S. cerevisiae* **a**-factor are recognized by enzymes that proteolytically remove the -AAX residues (Ste24 and Rce1), and then carboxymethylate the now C-terminal cysteine (Ste14) [5]. To begin, we consolidated the CAAX motifs of known pheromones in the Ascomycota into a dictionary to identify potential pheromone candidate ORFs assuming strong conservation of the motif as a very stringent cutoff (Supp Figure 1). To avoid biases based on the small number of known pheromones, we relaxed the CAAX detection dictionary based on studies of the possible combinatorial signatures of farnesylation, proteolysis and carboxymethylation in *S. cerevisiae* [21–23] (Supp Figure 1).

We then selected the list of ORFs ending in CAAX for candidates that have an in-frame methionine within 100 residues upstream of the stop codon to find ORF candidates that are within the expected size range of pheromone genes in Ascomycota (Figure 2A). In short, our pipeline considers genome assemblies contig-by-contig and assumes that the pheromones can be positioned anywhere. Since our algorithm relies on tracing the ORF starting from the Stop codon, it fails to locate many pheromone genes that contain introns, like those in *S. pombe*. We believe this is a reasonable simplification of the algorithm given the lower frequency of introns in Hemiascomycota yeasts (2-14% of all genes), relative to other eukaryotes [24].

**Figure 2.**
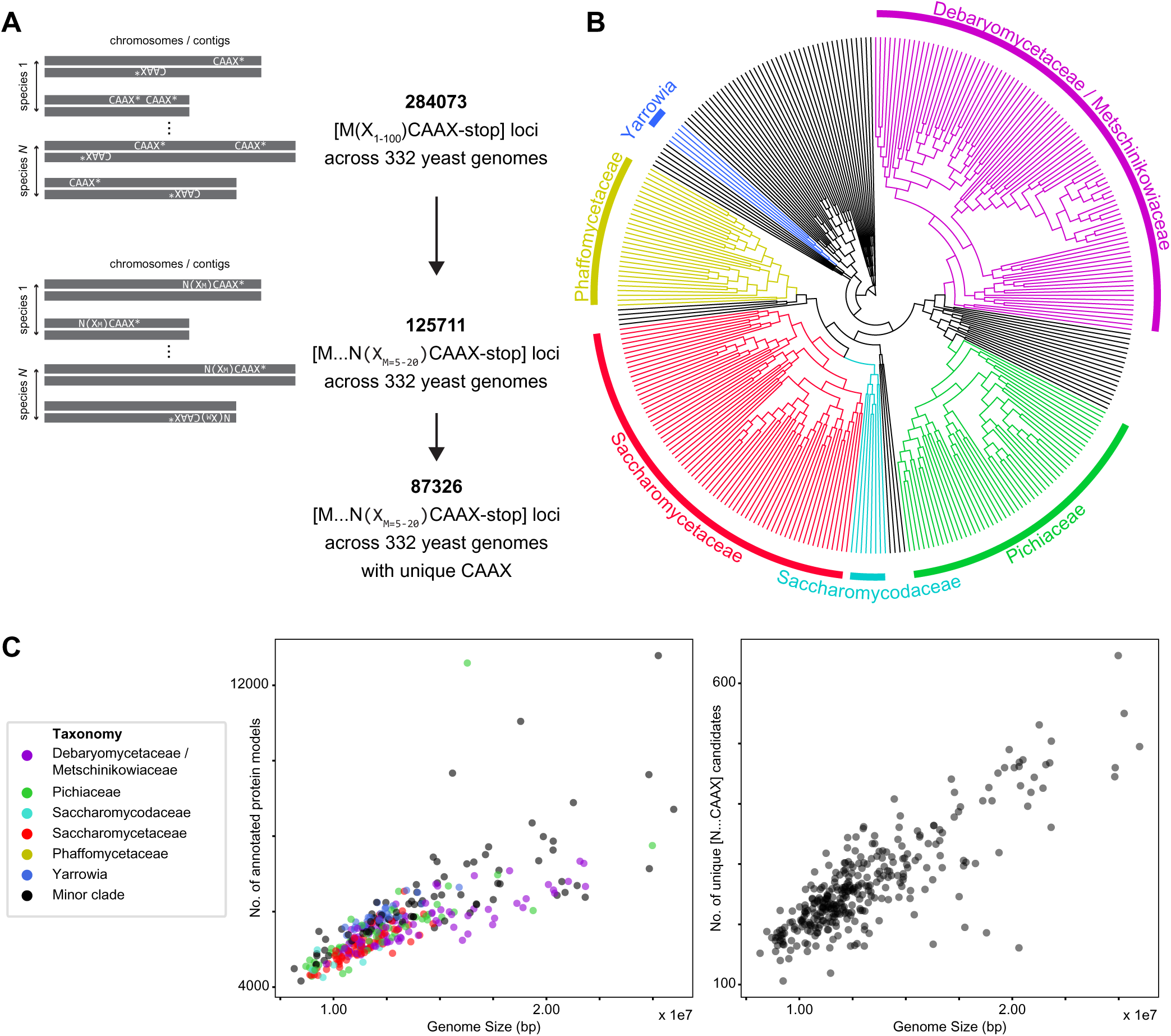
Fungal pheromone candidates can be identified by a progressive filter for small open-reading frames that are farnesylated and cleaved by a protease associated with mating. (**A**) Our algorithm begins by looking for all possible short open-reading frames with C-terminal farnesylation (CAAX-stop) in the 332 available fungal genomes, resulting in 284 073 candidates. We filter again for an in-frame proteolytic-motif asparagine (N) important for the final step of maturation to produce bio-active pheromone. This resulted in 125 711 candidates, encoded in 87 326 unique farnesylated loci. (**B**) Phylogenetic tree of all 332 sequenced yeasts across Saccharomycotina, covering clades like Debaryomycetaceae/Metschinikowiaceae, Pichiaceae, Saccharomycodaceae, Saccharomycetaceae, Phaffomycetaceae and Yarrowia that contain species important for both basic biology and industrial production. The tree is based on these yeast genome sequences with estimated relative divergence times. Selected clades are labeled. (C) On the left, a scatter plot shows strong correlation the number of annotated protein-coding genes with genome size, which ranges between 8 and 27 Mbp for the 332 yeast genomes. On the right, a scatter plot shows that the number of unique pheromone candidate loci is linearly correlated with genome size, ranging from 100 to 650 candidates, as expected of a greedy search for all possible pheromone candidates.

We ran this pipeline on a set of 332 published Saccharomycotina genomes [25] (Figure 2B and Table 1) and found 500-2000 candidate ORFs per genome, which is far larger than can be experimentally validated even for a single species. Subsequent to C-terminal farnesylation and carboxymethylation, *Sc***a**-factor precursors undergo N-terminal processing in two steps [5, 19]. The second proteolysis is the final step in maturation and is mediated by one of two Zn-dependent metalloendopeptidases (Axl1 and Ste23; Figure 1C) [26]. Random mutagenesis in *S. cerevisiae* **a**-factor revealed that the two residues immediately before and after the cleavage site are important for *Sc*Axl1 cleavage [19]. Comparing known Ascomycota pheromones, we noticed that the cleavage site is typified by N-X with X representing aromatic or uncharged residues and the peptidase cleaving C-terminal of the asparagine (Figure 1C and Supp. Figure 1A). Therefore, we required pheromone candidates to have an asparagine in frame with the conserved cysteine such that the mature pheromone is between 5-20 residues from residue X following the cleavage site N to the farnesylated C. Adding this sieve for an internal cleavage site between 5-20 residues from CAAX reduces the number of unique pheromone candidates to 100-700 per genome (Figure 2C). As expected, the distribution of pheromone candidates per genome scales with genome size similar to the number of annotated protein models (Figure 2C). Candidates are still too numerous for experimental validation, but further knowledge of pheromone biology can be used to identify the best candidates.

**Table 1.**
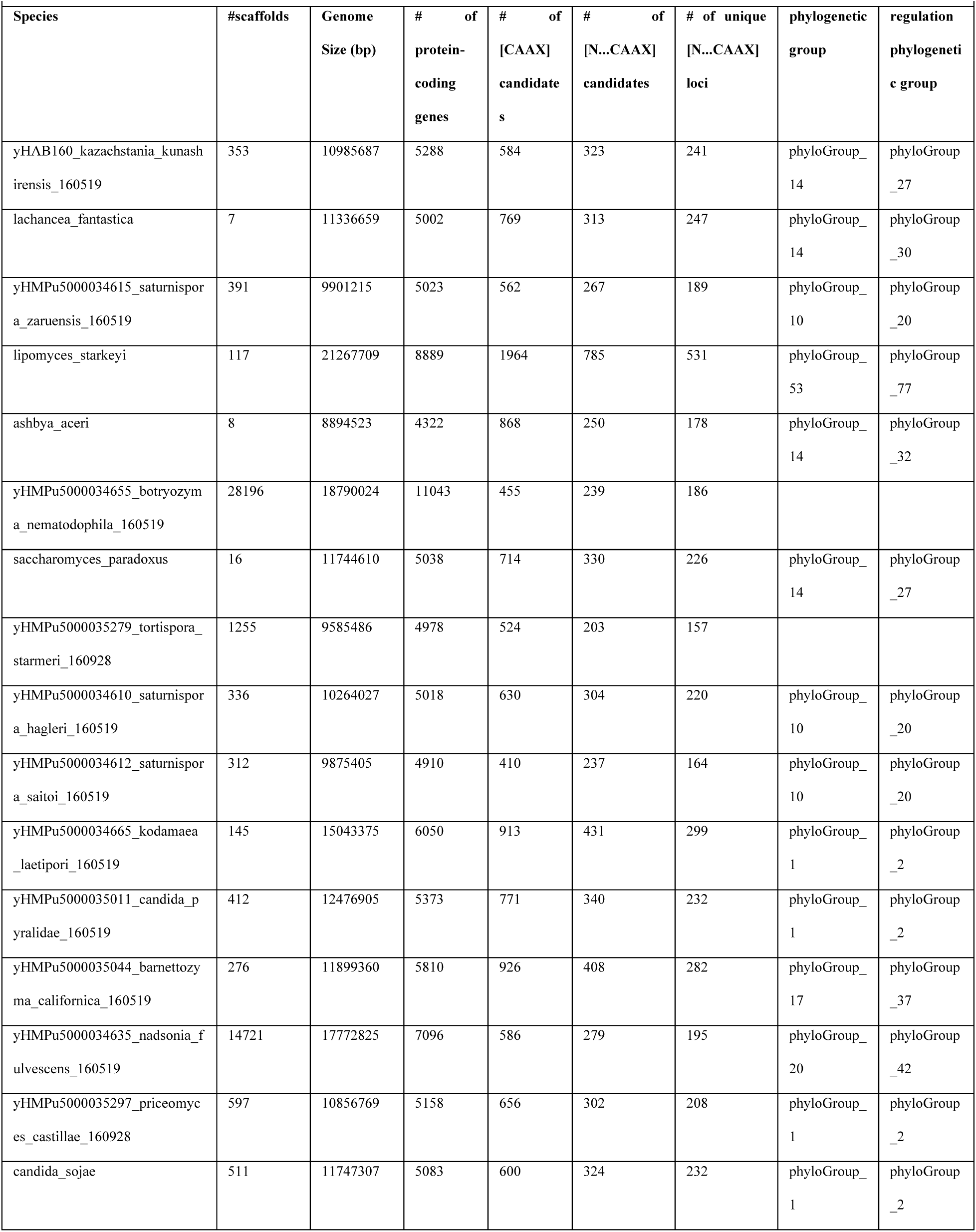
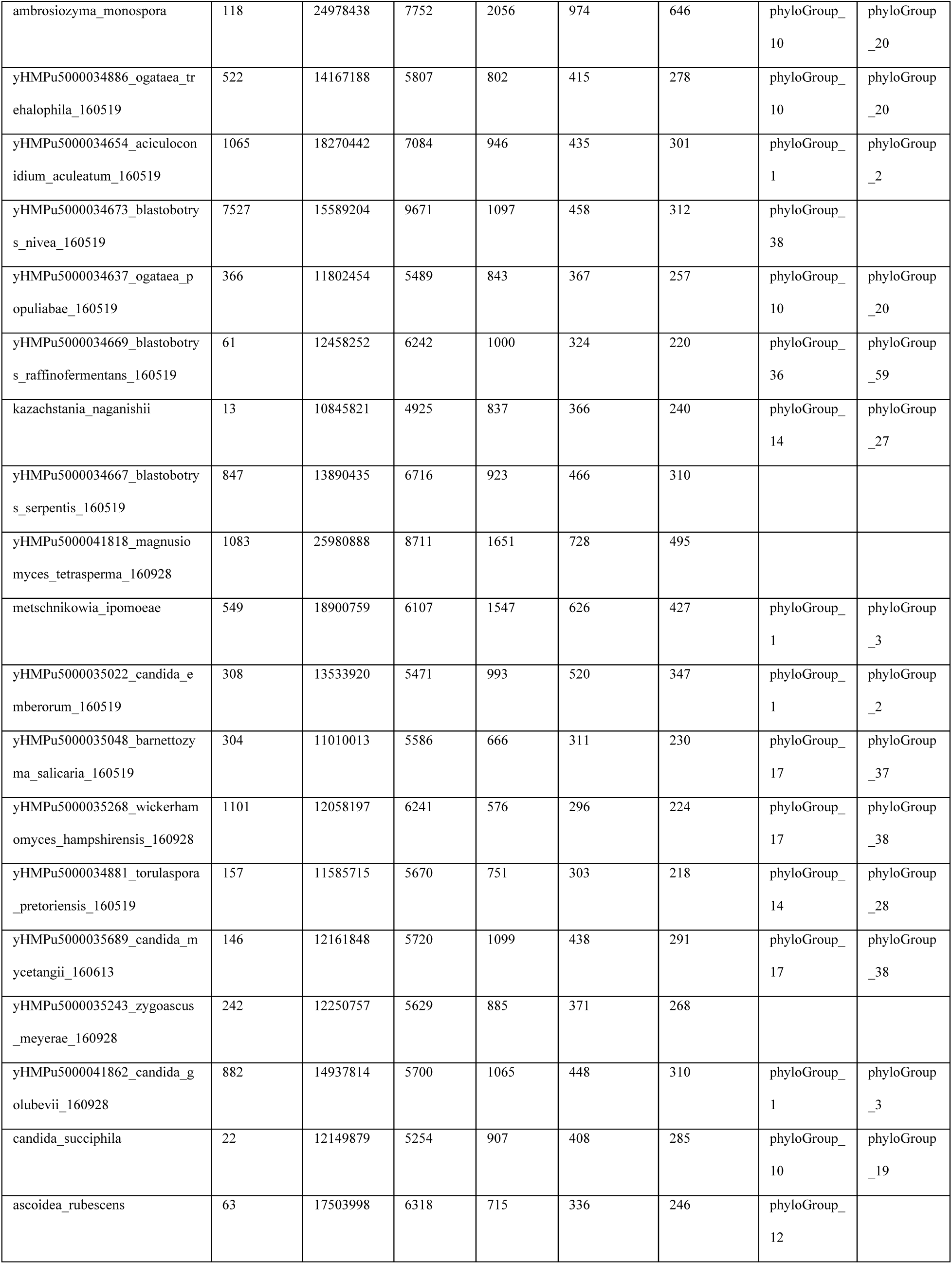
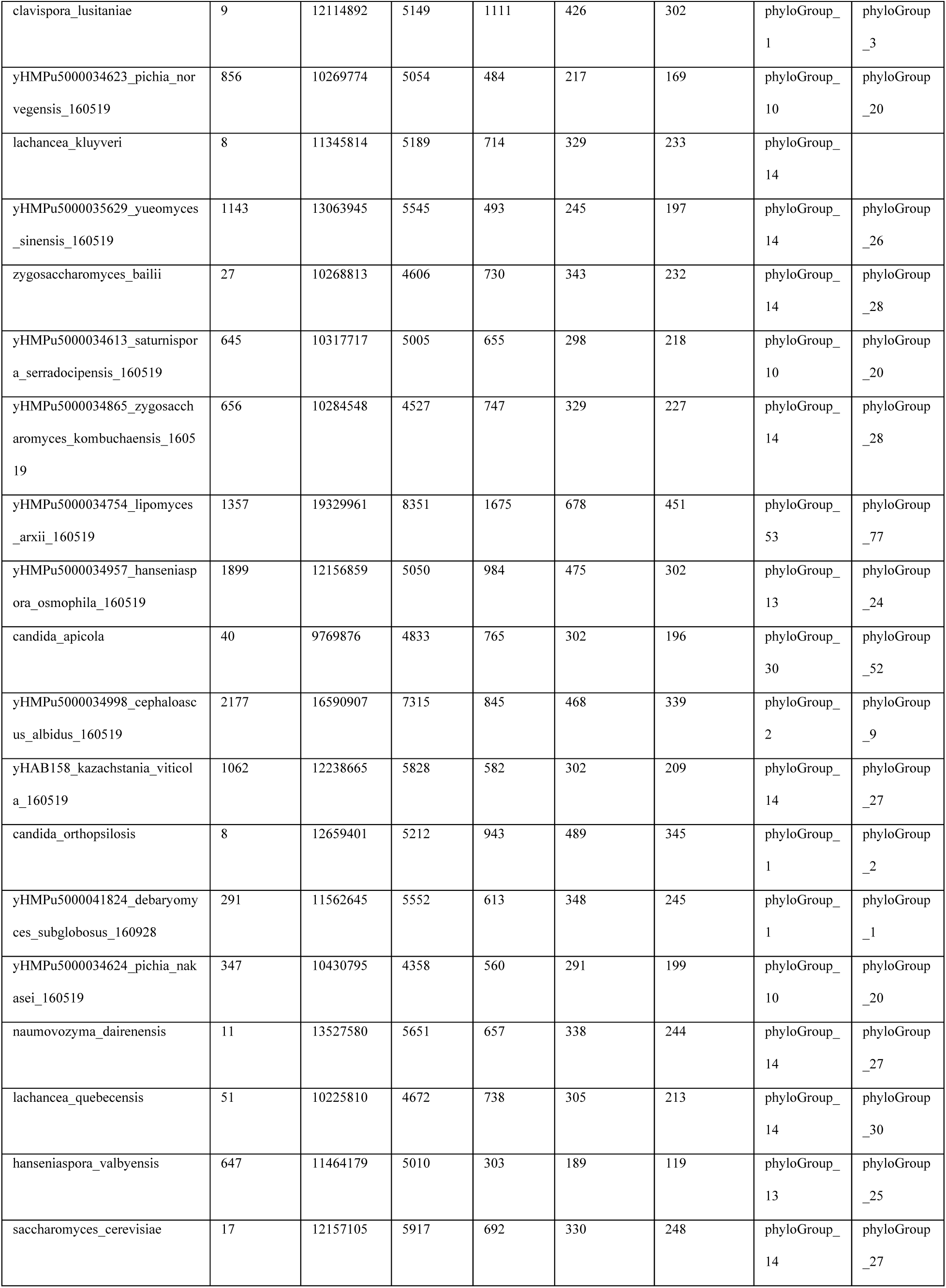
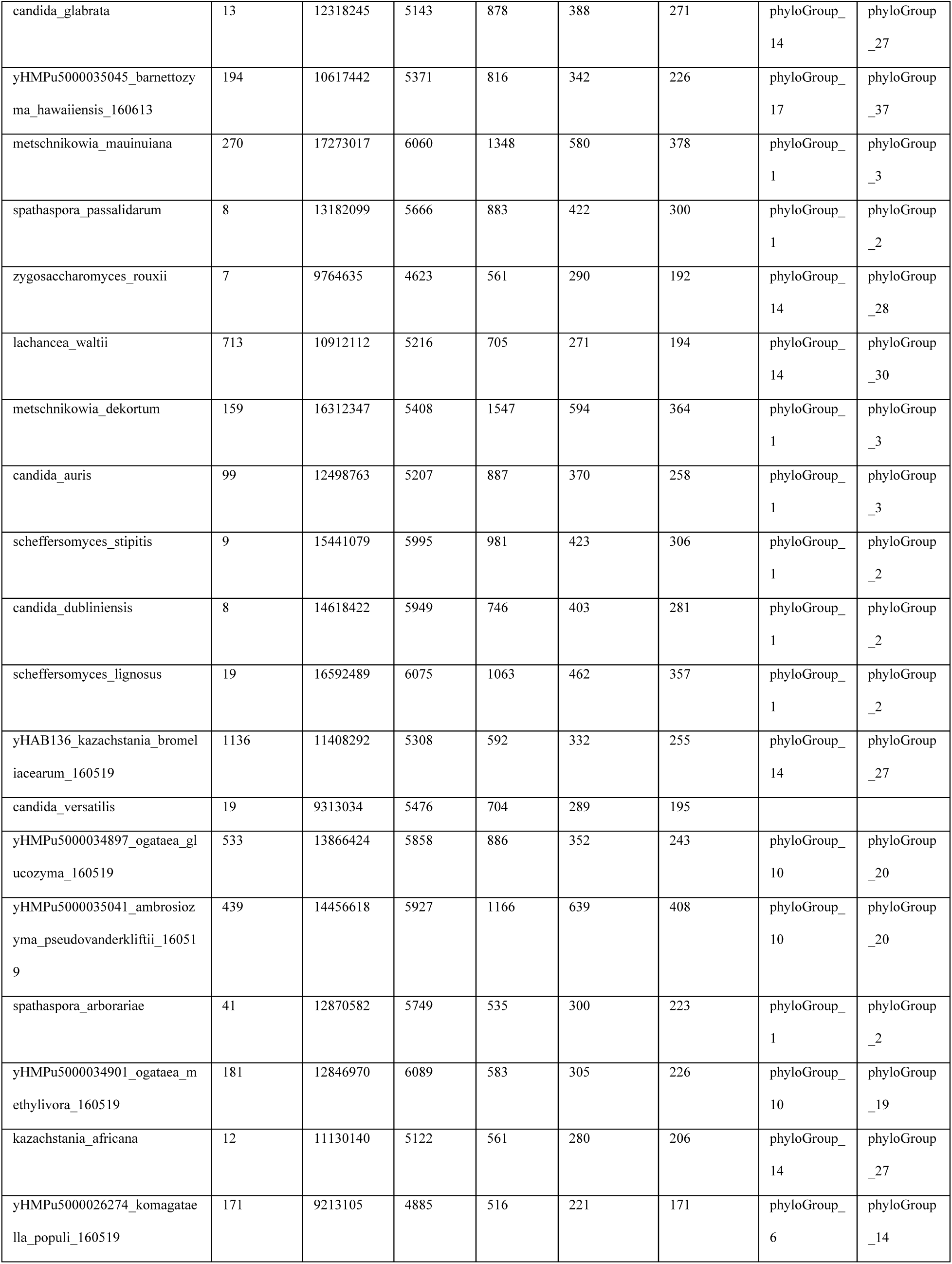
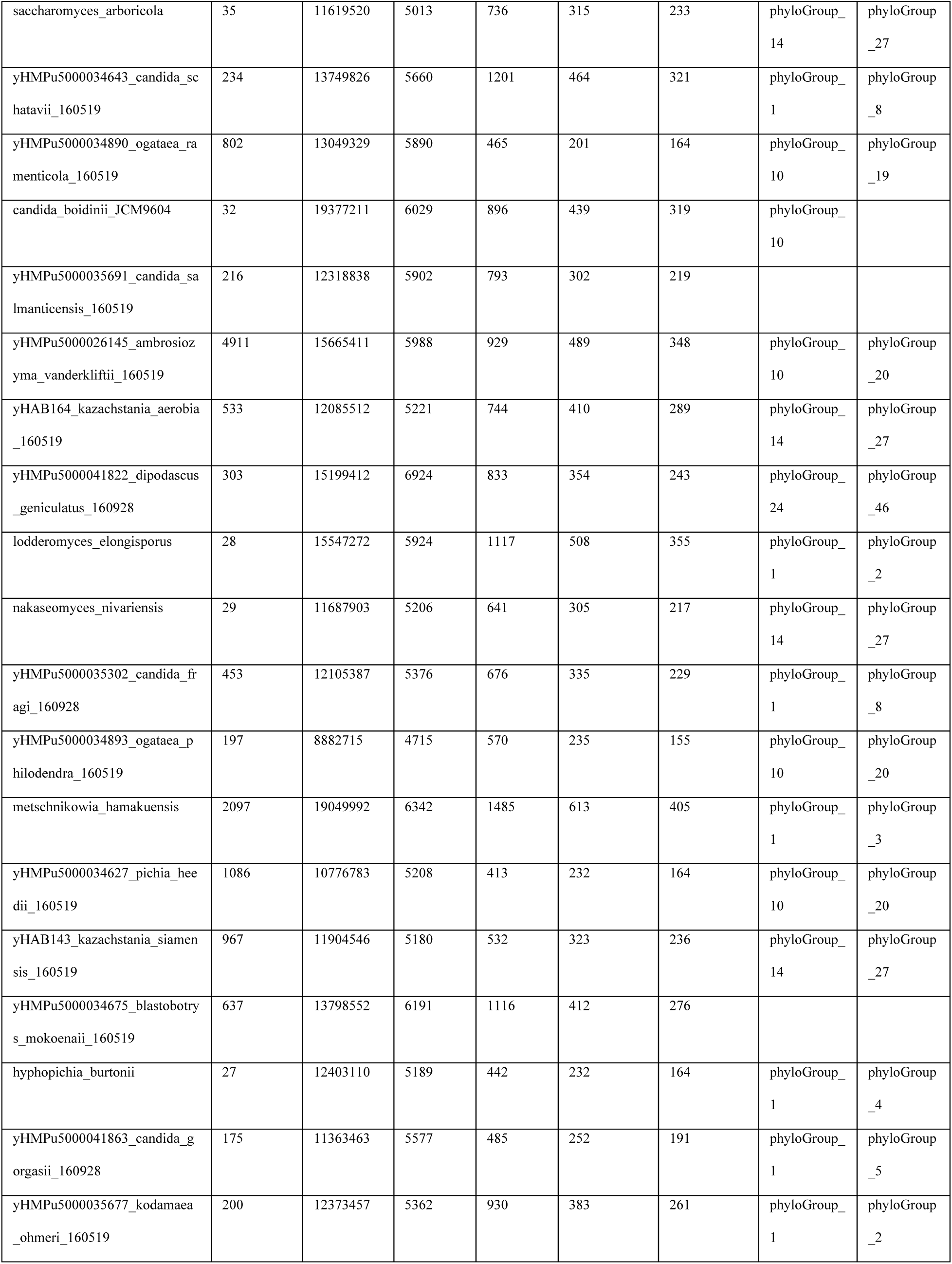
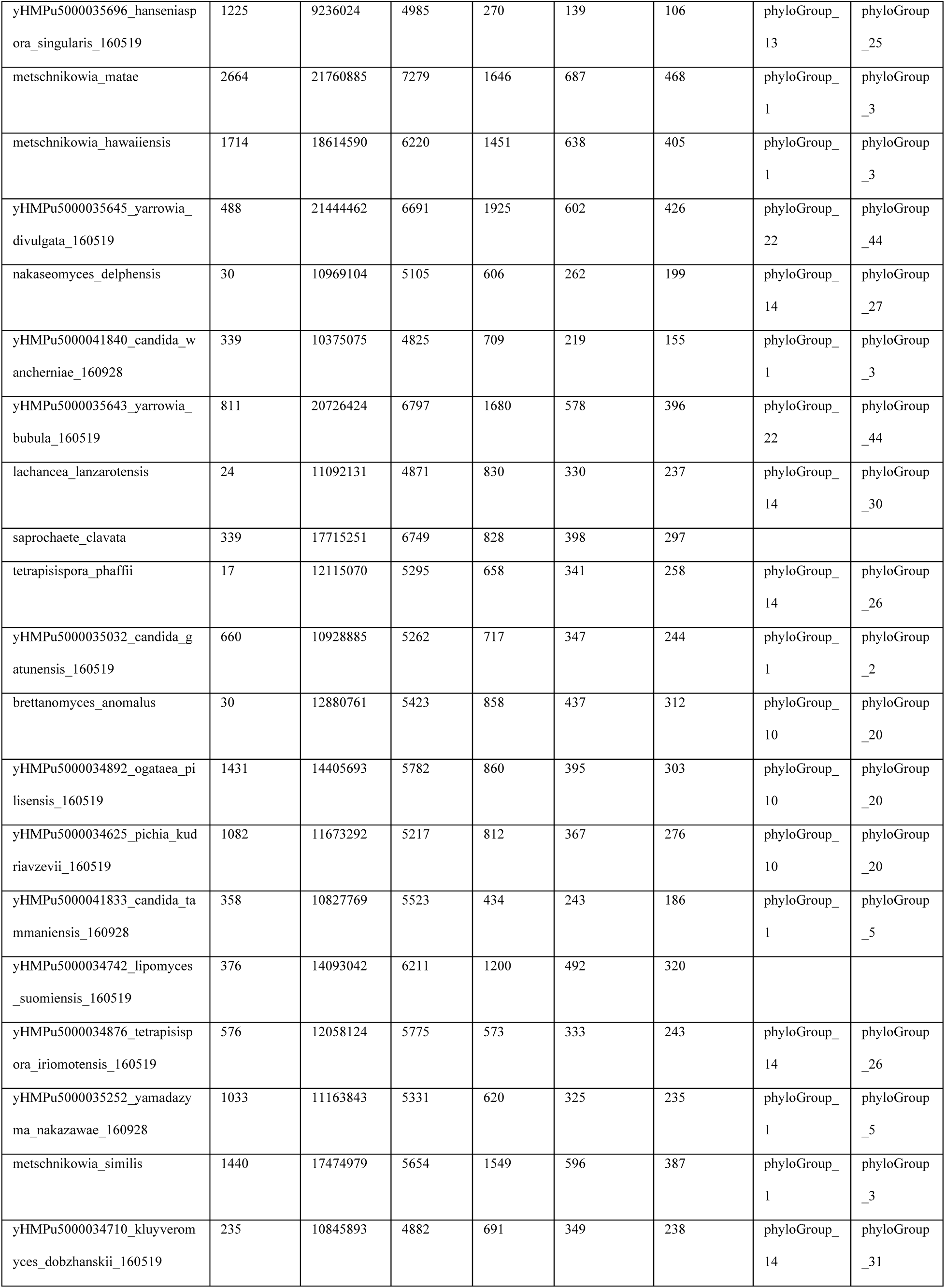
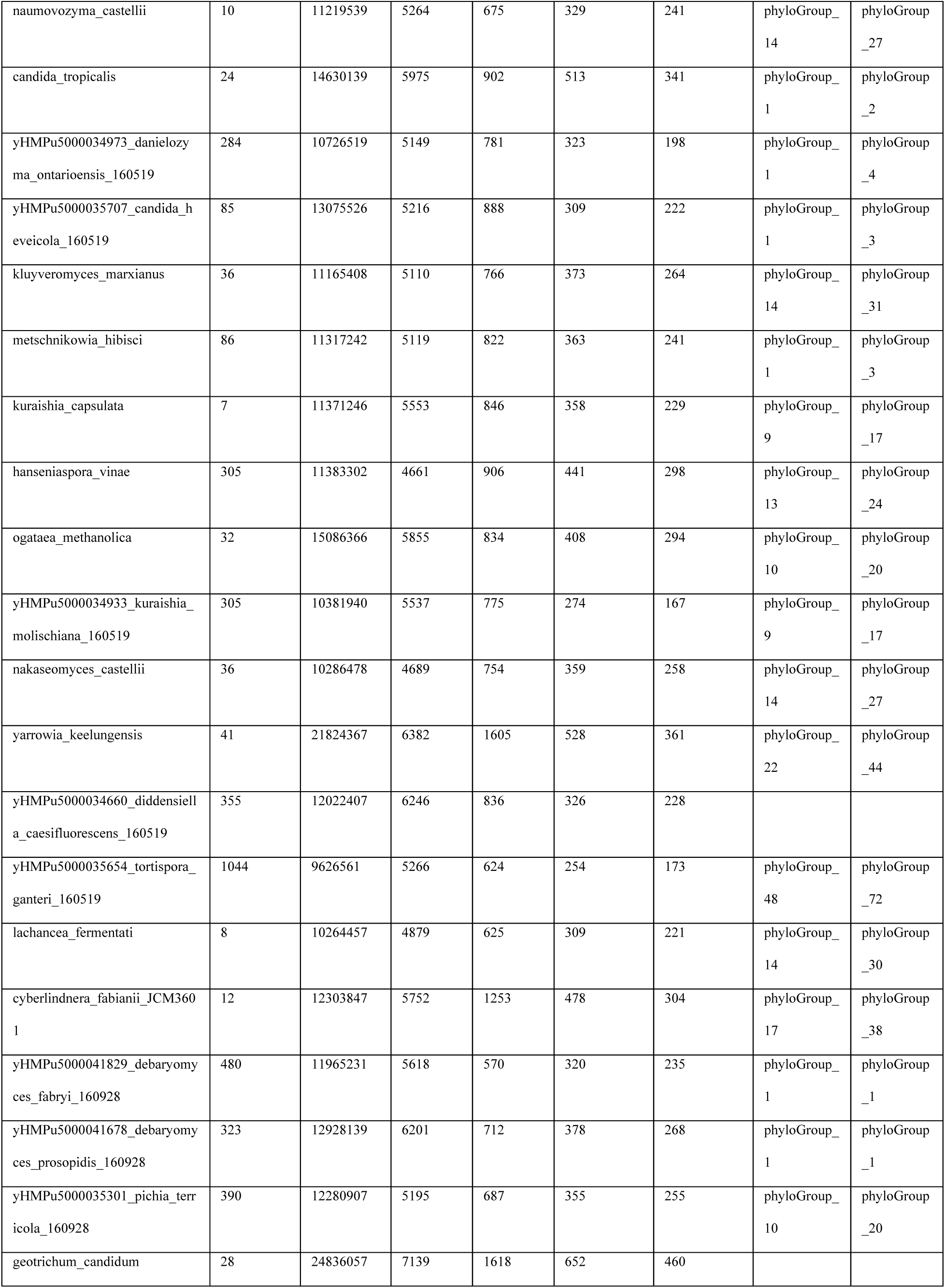
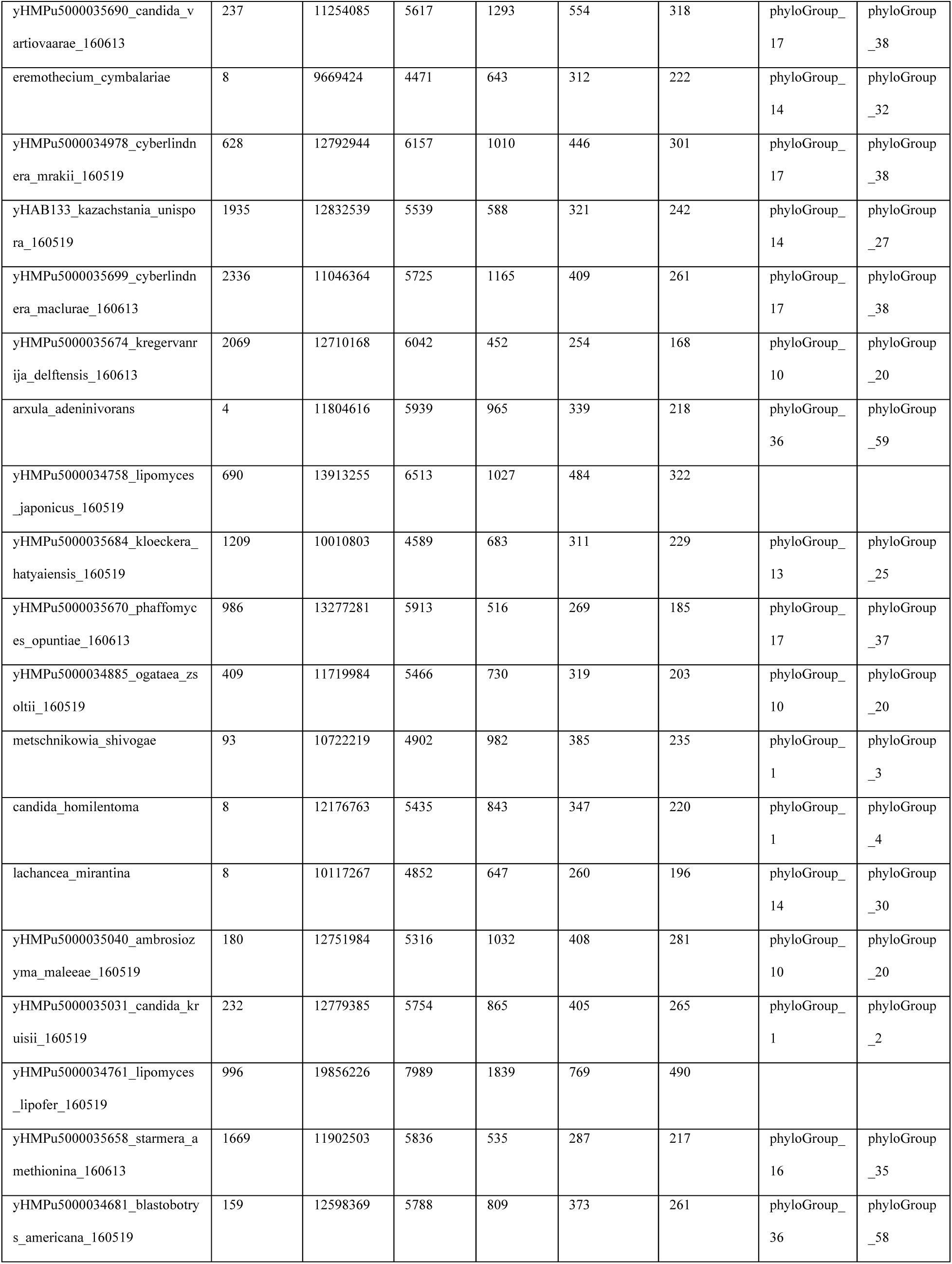
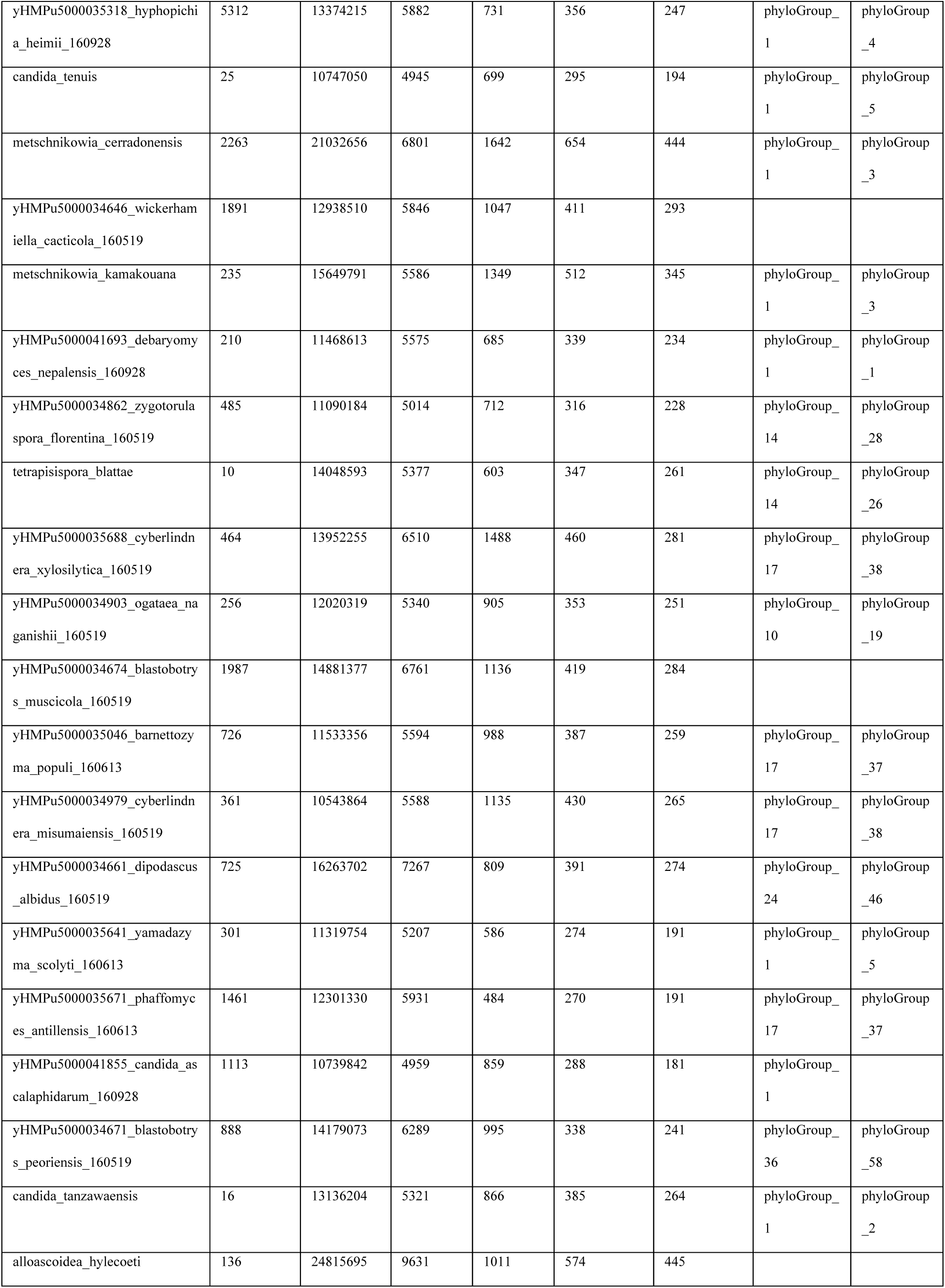
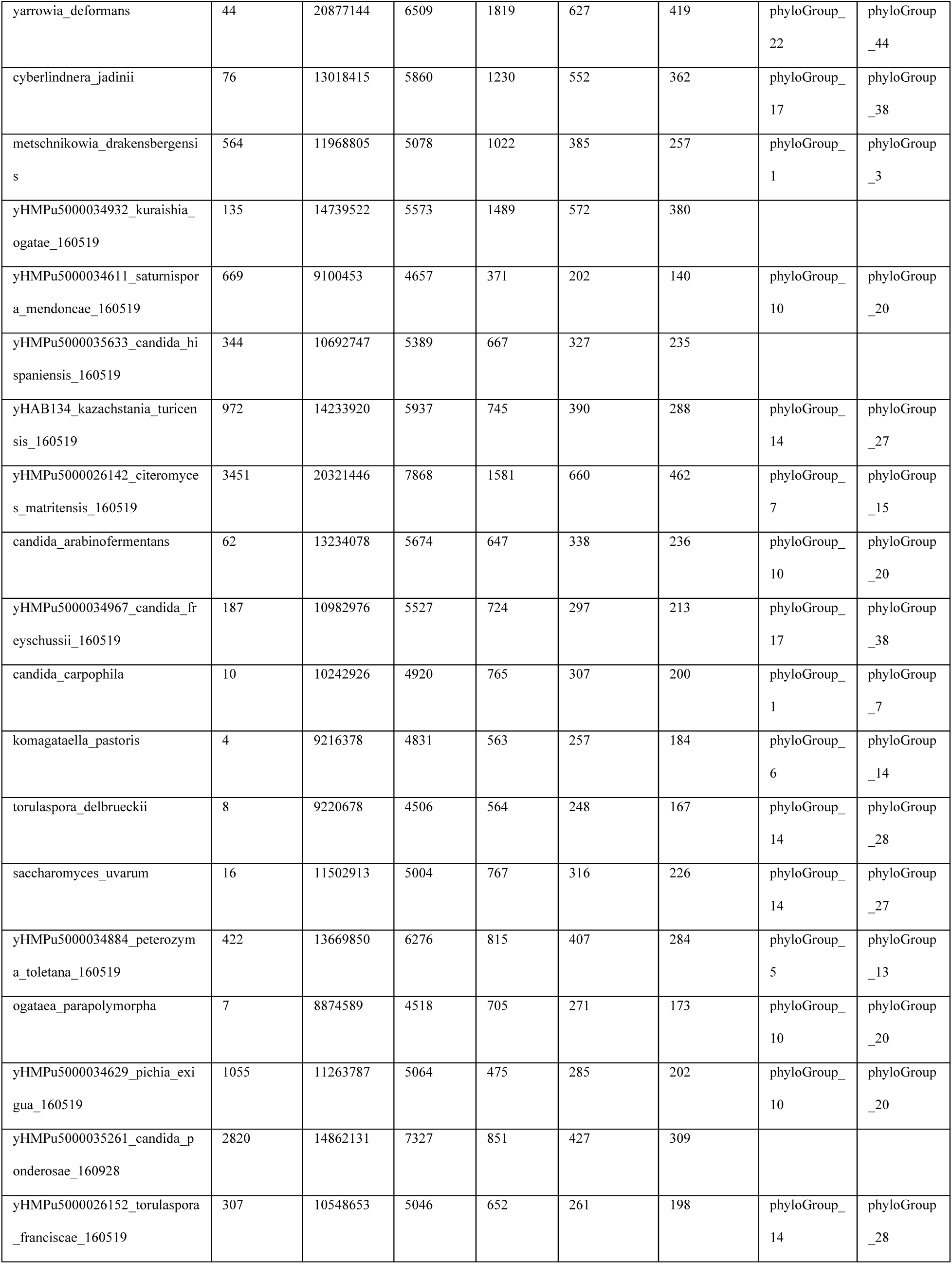
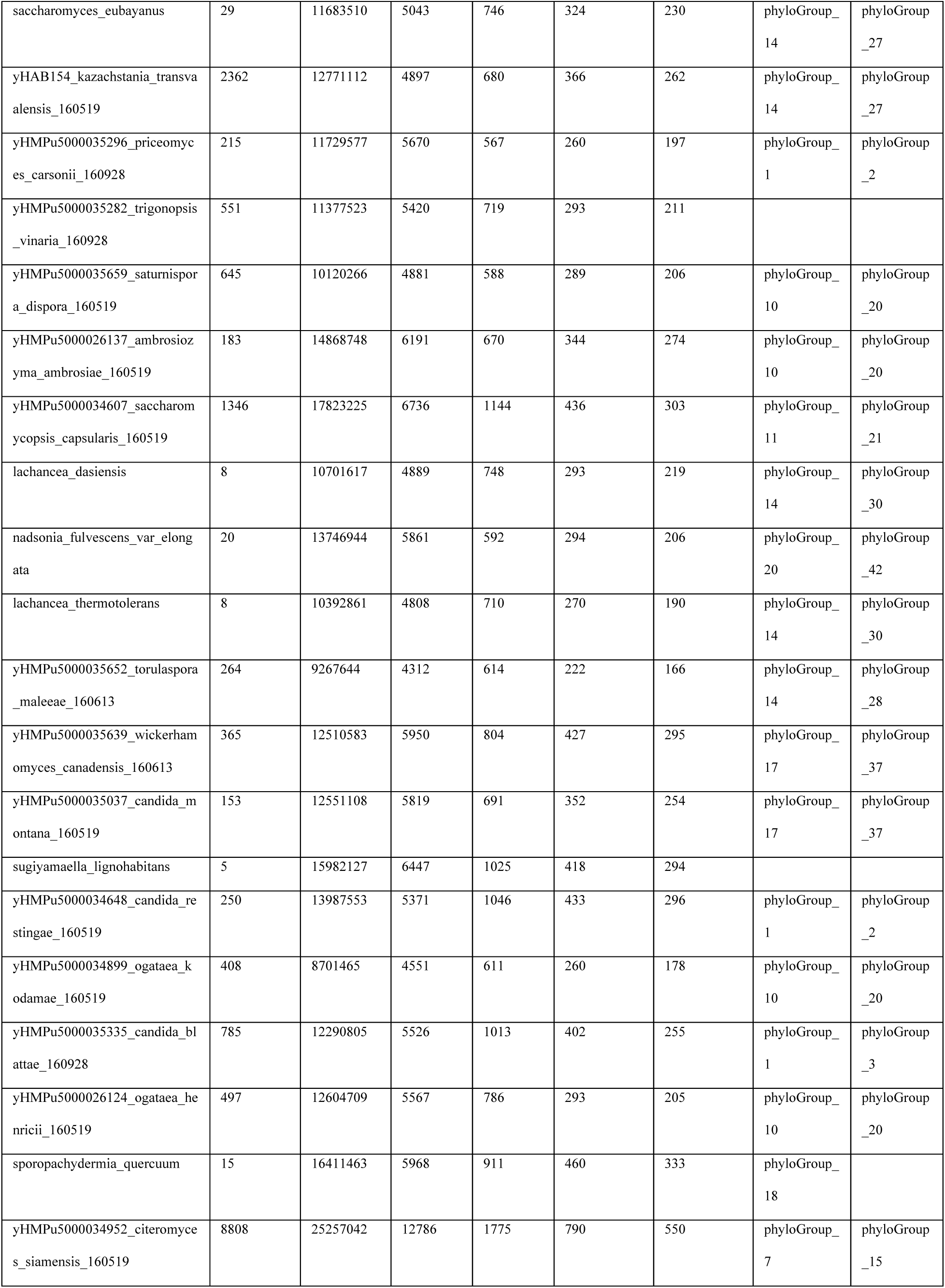
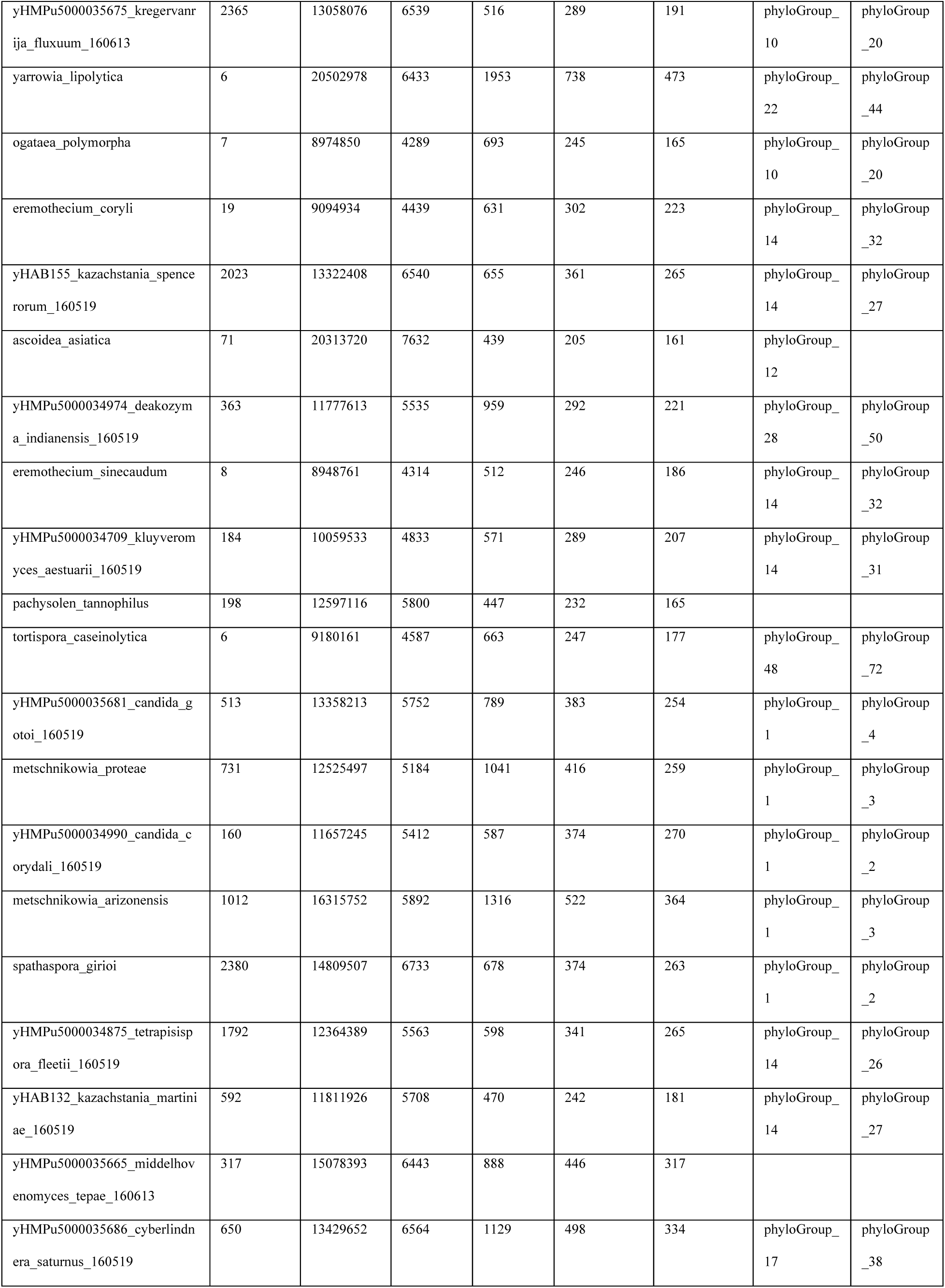
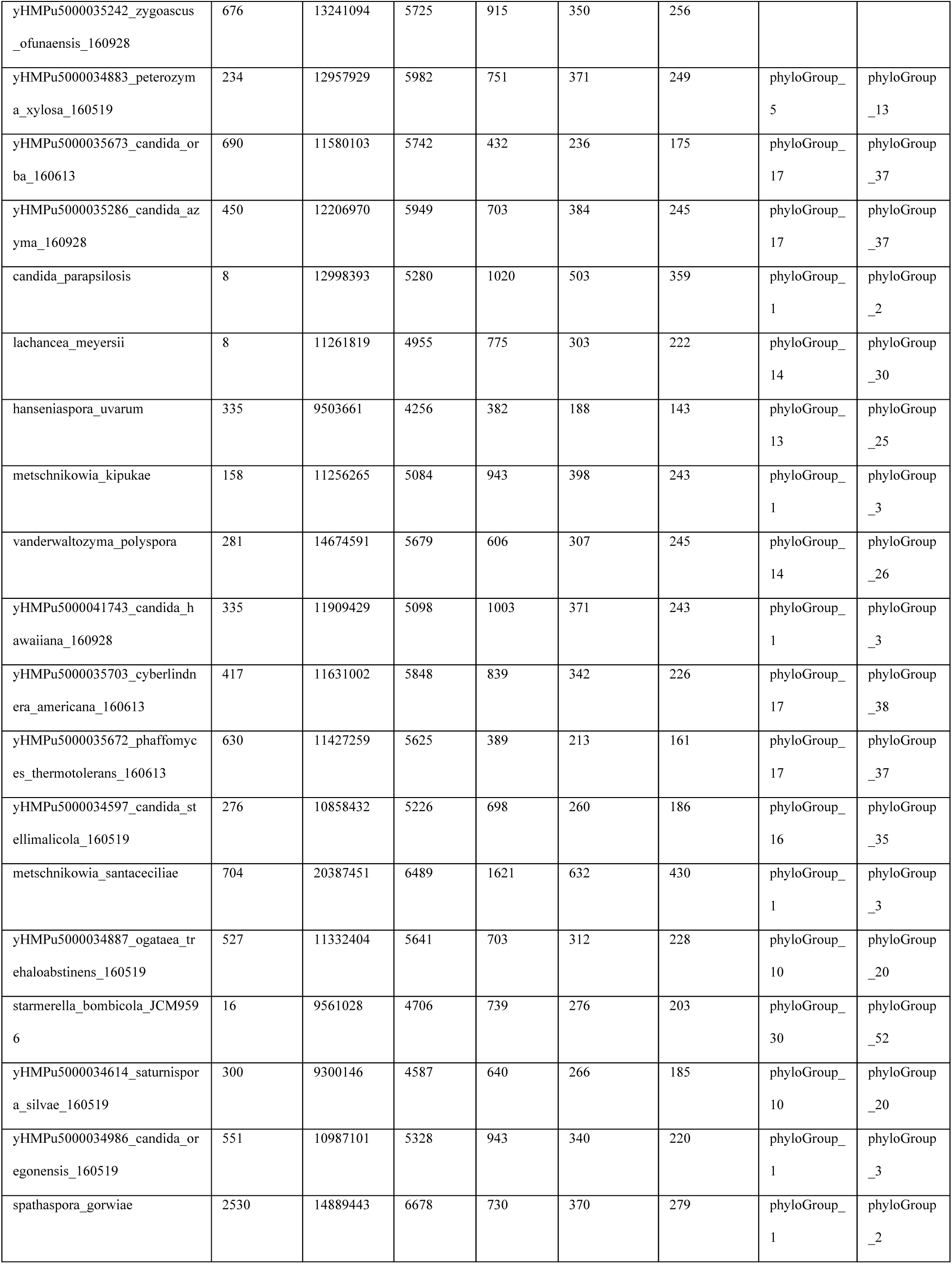
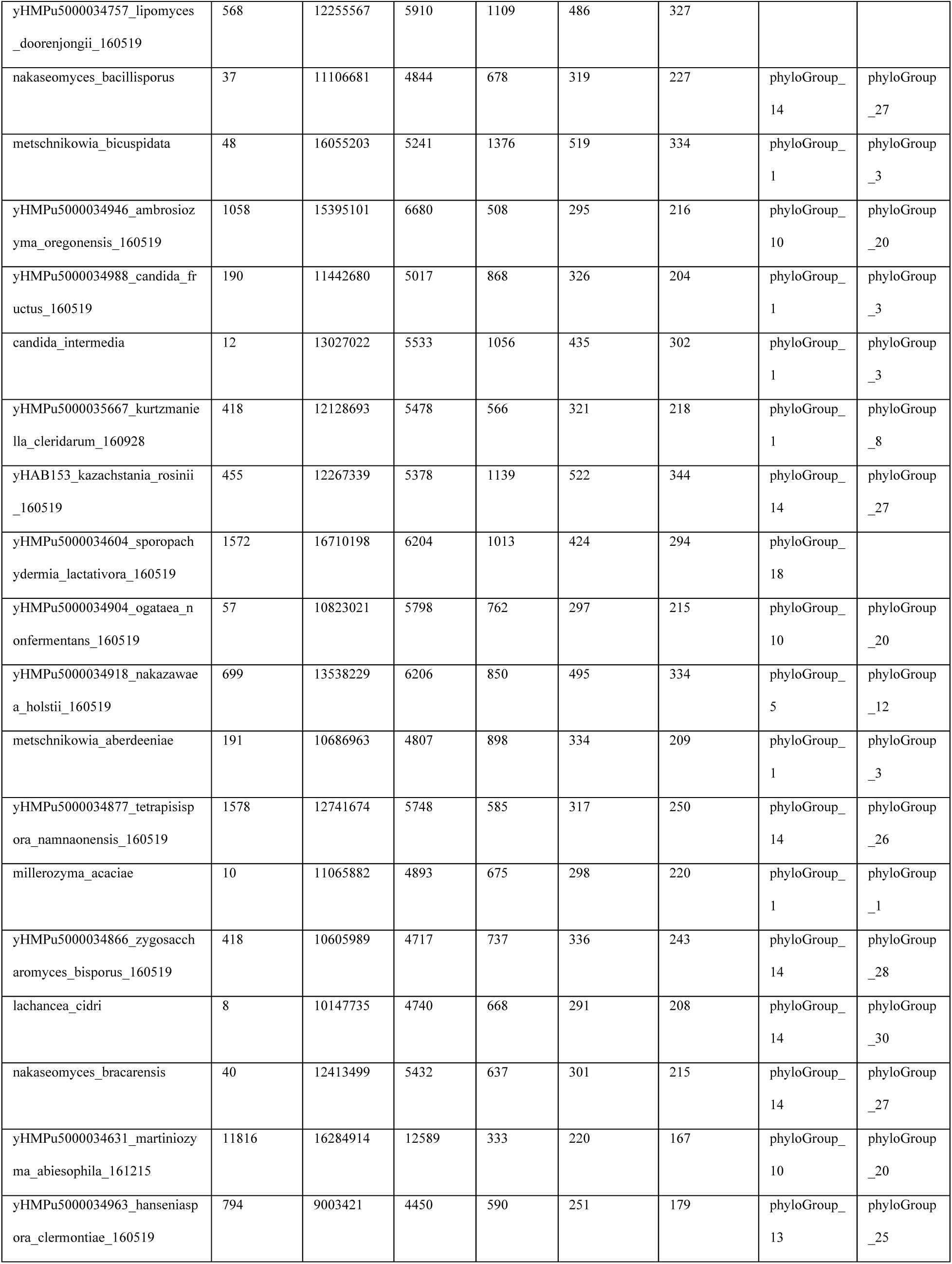
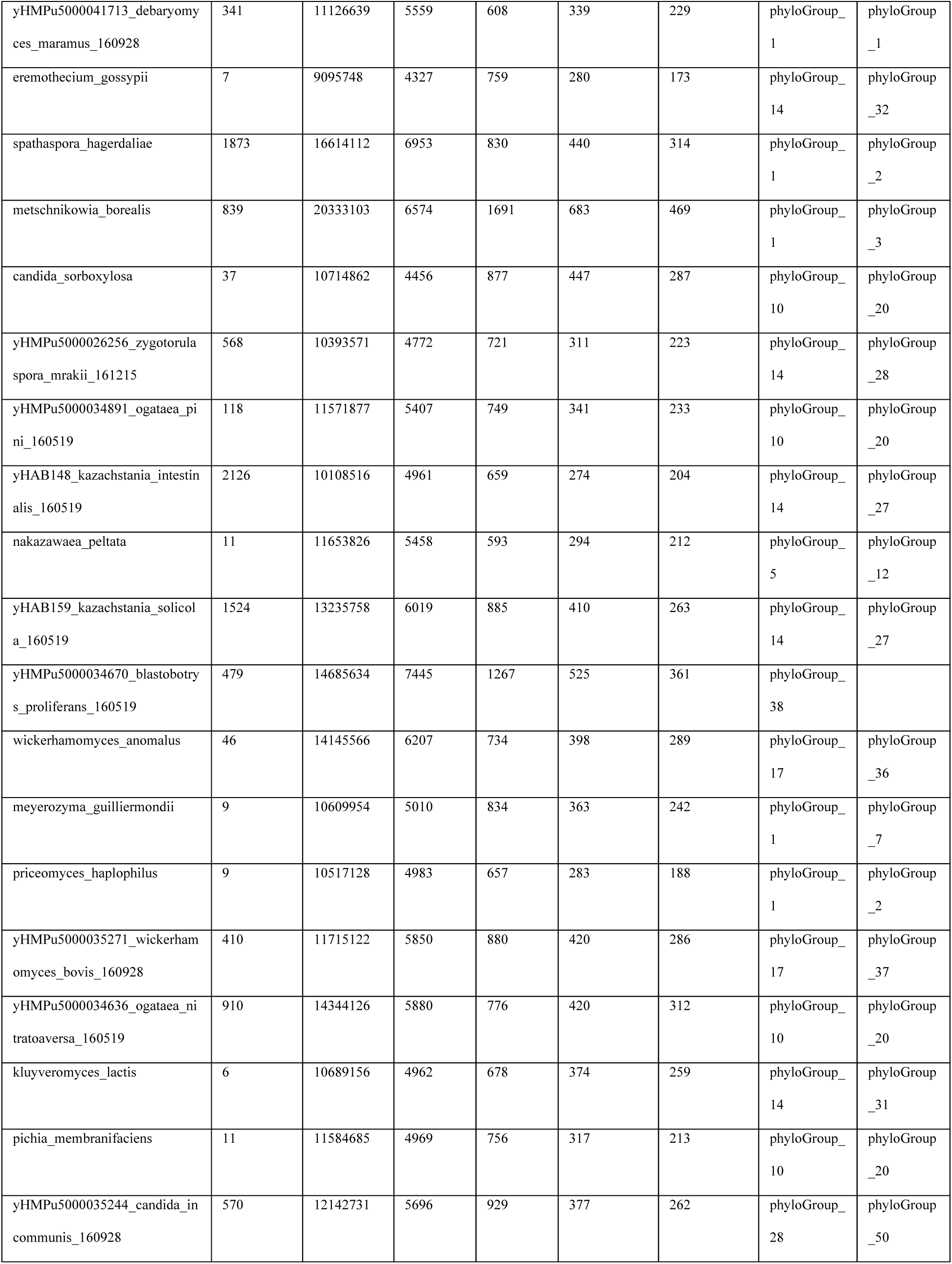
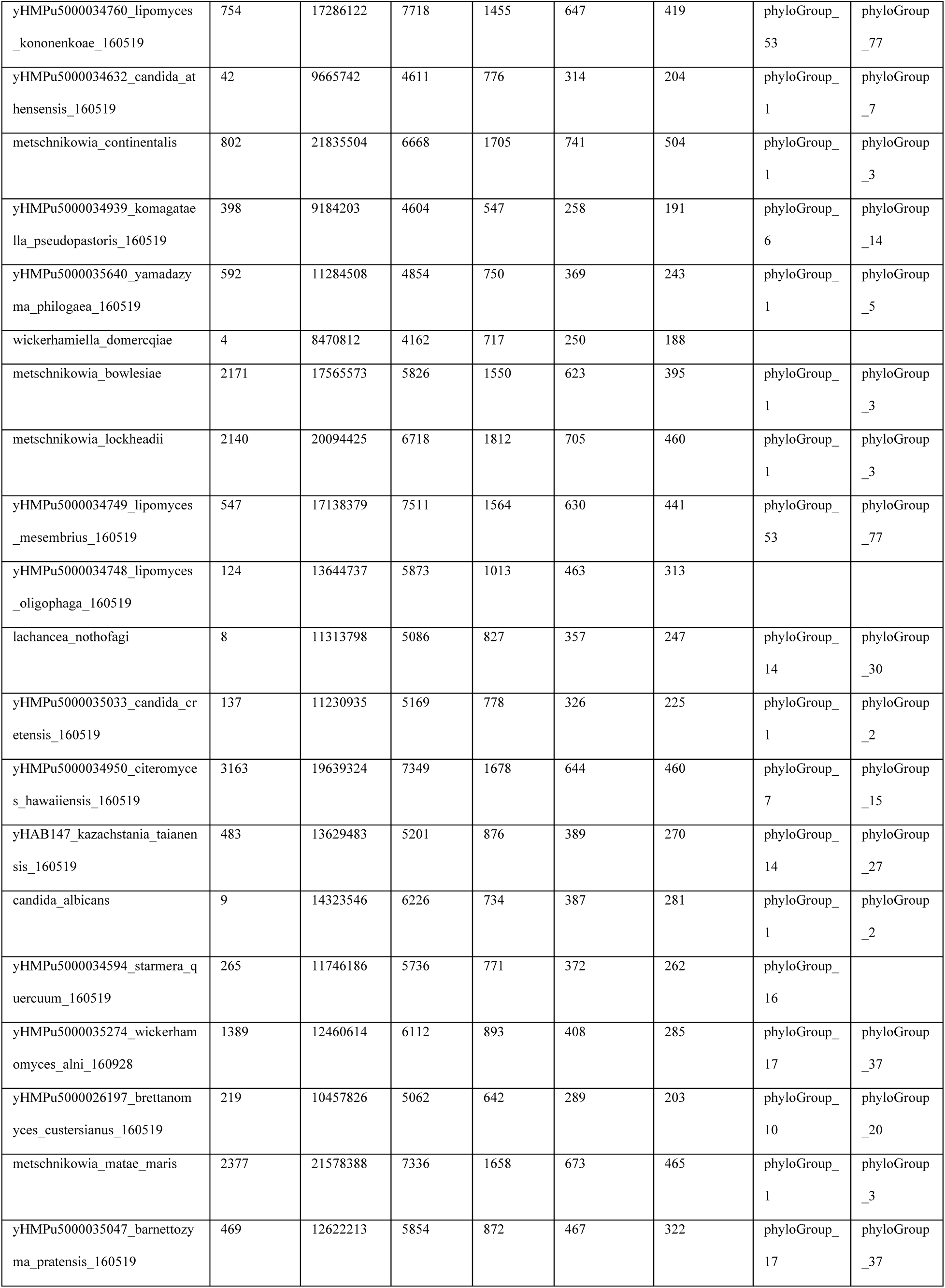
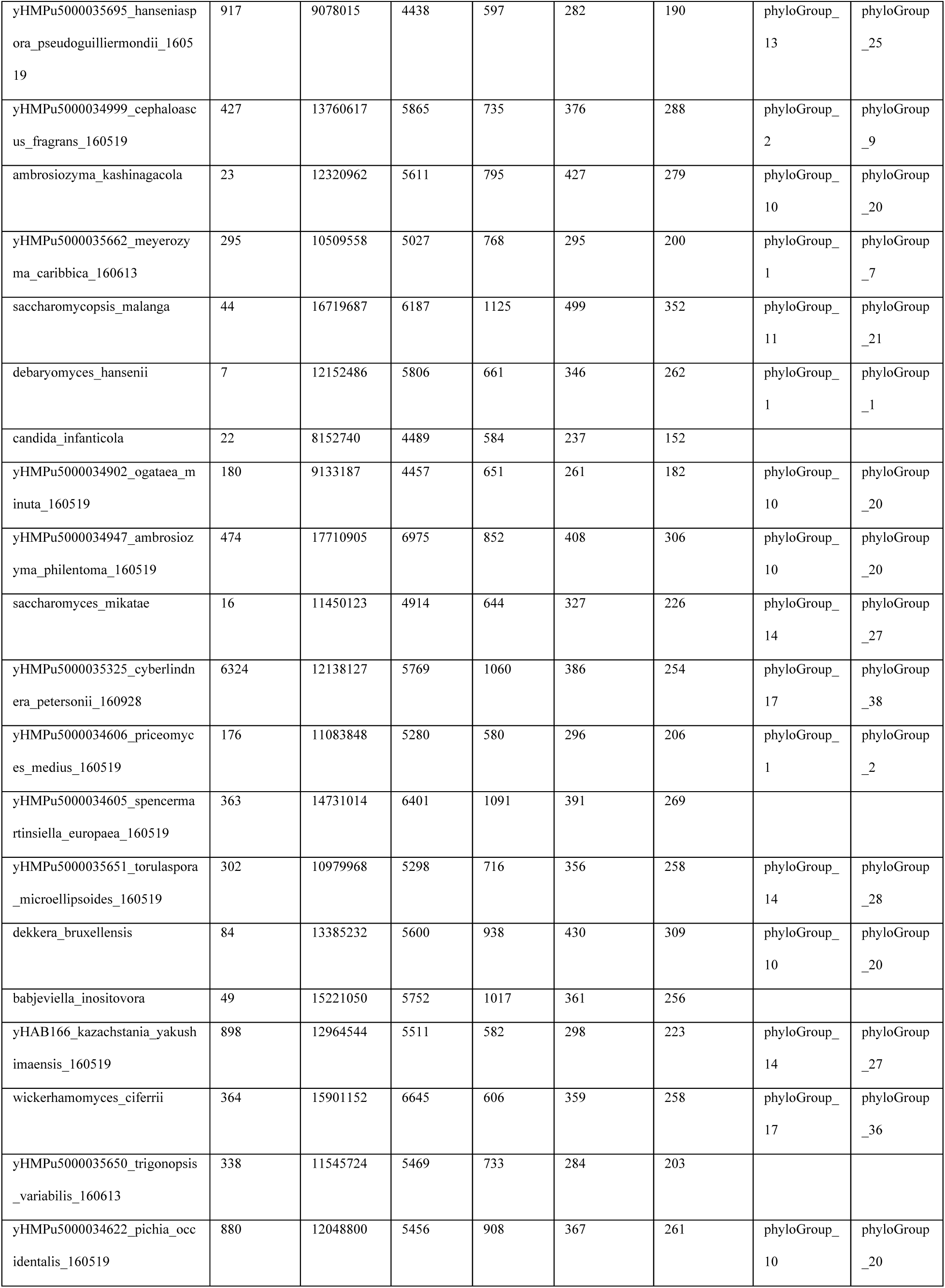
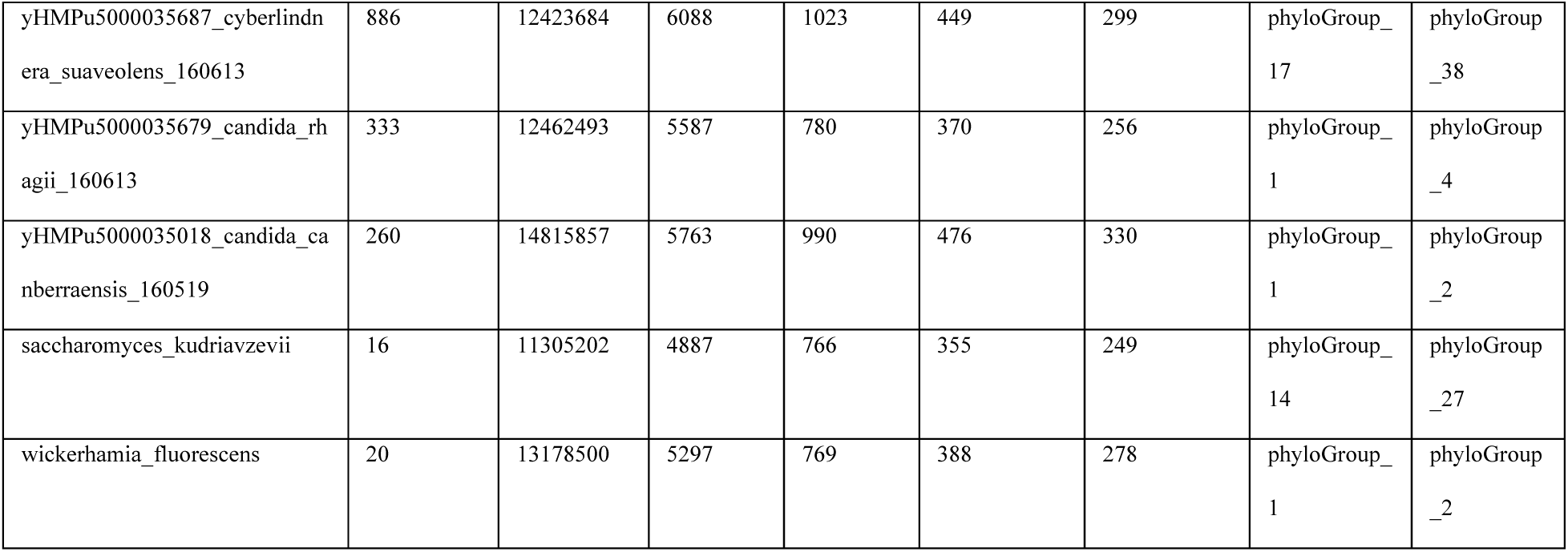
Results of computational filter on 332 yeast genomes.

### Yeast pheromones are encoded in multiple gene copies and are conserved among closely related species

Yeast pheromones of closely related species are quite similar, although the number of copies in each genome varies (e.g. 1 each in *Candida albicans* and *C. dubliensis* [27], 2 each in *Komagataella phaffi* and *K. pastoris* [28], and 1 to 3 in different *Saccharomycetaceae* [18]). The amino acid sequence of the ORFs is most strongly conserved within the mature pheromone peptide sequence which must interact both with the **a**-factor exporter, Ste6, and the **a**-factor receptor, Ste2. The *Saccharomycetaceae* is the largest yeast lineage across which a pheromone candidate is known to be conserved and we used its time of divergence as an evolutionary horizon within which species are likely to show strong pheromone homology (Figure 3A). We identified all monophyletic clades on the yeast phylogram with ancestral nodes within this time horizon: 300 species give rise to 23 phylogroups, each containing at least 2 species, and 32 species are singleton orphans with no related species within this horizon (Figure 3A, B). Among these 23 phylogroups are recognized clades of yeasts, including *Debaryomycetaceae/Metschinikowiaceae, Pichiaceae*, *Saccharomycodaceae*, *Saccharomycetaceae*, *Phaffomycetaceae* and *Yarrowia*, which contain well-known species (Figure 3B). By pooling the candidates of species within these groups, we expect that the best candidate pheromones will share two properties: they are strongly conserved and conservation is strongest in the mature region, between the predicted N- and C-terminal processing signals.

**Figure 3.**
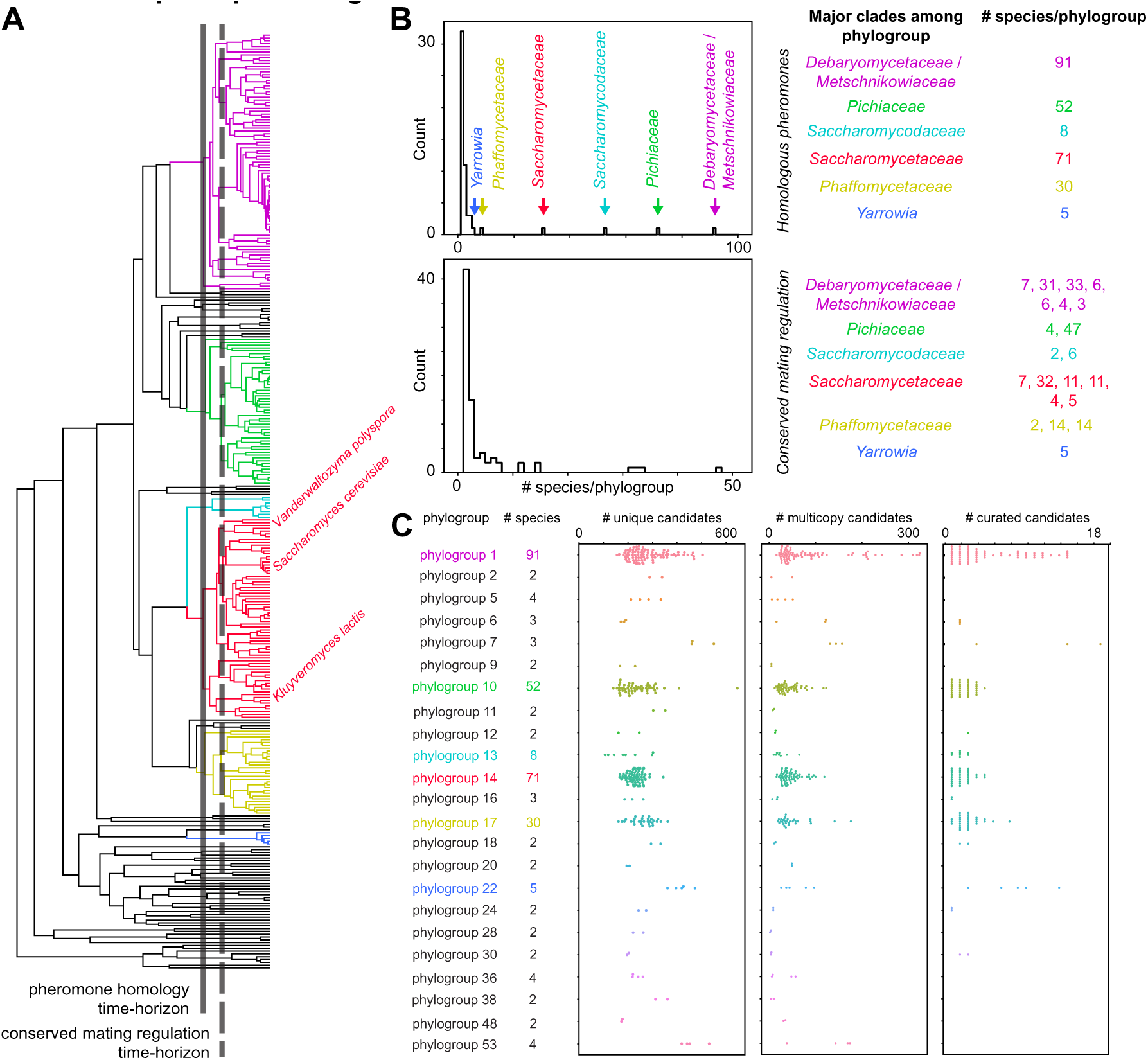
Fungal pheromones are conserved within closely related species and are often encoded in multiple copies in a genome. (**A**) Phylogenetic tree describing the evolutionary relationship of 332 yeasts with two horizons indicated. The solid black line indicates the maximum evolutionary distance at which mature pheromone sequences show detectable sequence homology, corresponding to the divergence time of *S. cerevisiae* and *Kluveromyces lactis*. The dashed black line indicates the maximum evolutionary distance at which mating genes share conserved regulatory motifs, corresponding to the divergence time of *S. cerevisiae* and *Vanderwaltozyma polyspora*. (**B**) Based on the pheromone homology time horizon (top), we separated 332 yeast genomes into **23** phylogroups of at least 2 species and 32 singletons. The most populated phylogroups correspond to the listed well-known clades where closely related species have been densely sequenced. These clades are represented in the tree by the corresponding colors. Phylogroups expected to share mating regulation (bottom) are smaller with the larger clades divided between multiple phylogroups as indicated; 290 genomes are distributed across 37 mating regulation phylogroups of at least 2 species with 42 singletons. (**C**) Number of pheromone candidates per species plotted for each of the 23 conserved-pheromone phylogroups (different colored circle swarms), where each circle corresponds to the number of candidates in a species, and each copy of a group of closely homologous sequences within a genome is counted separately. Selecting for candidates that have at least two homologous copies within the clade reduces the number of viable candidates per genome to 10-300 (center compared to left). Manual curation of candidates similar to known pheromones identifies the most likely pheromone(s) in each genome (right from center) for experimental validation. There are 1-19 curated candidates encoded in each species for experimental testing. Some phylogroups contain too few species and no candidates rose above the rest through manual curation.

Farnesylated (**a**-factor-like) pheromones are often encoded in multiple copies in a yeast genome, presumably to secrete sufficient pheromone from non-motile, haploid cells. The copy number of the pheromone genes seems to vary rapidly even among closely related species [18, 29]. To identify the most likely pheromones, we looked for candidates that have multiple copies across genomes of at least some of the species in the clade (at least 2 homologous copies in the same genome, or homologous copies in 2 species of the same phylogroup). Since only the mature region of the pheromone is bioactive, this filter identifies gene copies based on the conservation of the amino-acid sequence between the predicted protease site (Asn) and the stop codon. Specifically, candidates that are greater than 85% identical in the mature region are considered copies of the same predicted pheromone in the genome. Our algorithm only considers ORFs that maintain the reading frame between the start and stop codons in a single exon and will not identify gene copies that contain introns or frameshifts.

Of the 78 206 candidate ORFs that have both farnesylation and proteolysis sites (N…CAAX candidates) across 300 species in the 23 phylogenetic clades, 11 065 candidates have at least one other homologous candidate in the phylogroup. There are between 10-300 candidate genes per species, with false positive ORFs that are conserved among closely related species present in a number of clades (Figure 3C, middle). Manual curation of the multicopy candidates by similarity of amino acid composition to known pheromones and presence across multiple species narrows down the candidates to a total of 705 loci across 218 species with an average of 3.2 ORFs that encode potential pheromones per genome, and a range of 1-19 candidates per genome (Figure 3). These 705 candidates mostly account for a “best” pheromone candidate often encoded in multiple copies per genome.

The well characterized farnesylated pheromones of Saccharomyces species [6, 18], *C. albicans* and *C. dubliensis* [27], and *K. pastoris* and *K. phaffi* [28] clades are identified among candidates with multiple copies by our algorithm (Supp. Figure 1 and Supp. Table 1). The Saccharomyces pheromone candidates are a large group with many homologous copies among the genomes of the parent clade of Saccharomycetaceae. The pheromones of *C. albicans* and *C. dubliensis* are homologous to each other and only encoded in one copy in each genome. The Komagatella pheromones are identified by the presence of the same pheromone in the sequenced sister species *K. pseudopastoris* and *K. populi*. Some known pheromone genes in *Saccharomycetaceae* identified by synteny are not >85% identical and were not identified in our filter. To account for slightly more divergent pheromones, we filtered for candidates that are >70% identical (17 125 of 78 206 candidates), expanding the list to 792 candidates in 238 species (Figure 3C, right and Supp. Table 1). Even with this expanded list, some species in the large clades of *Debaryomycetaceae/Metschinikowiaceae* (21 of 91 species), *Pichiaceae* (11 of 52 species) *and Phaffomycetaceae* (1 of 30 species) are missing candidates that are homologous to the best candidates in the most closely related species (Supp Figure 3). Unfortunately, there are many explanations for this, including lineage-specific selection for faster pheromone evolution, incomplete or poorly assembled genomes, or the presence of introns in pheromone genes.

### Pheromone candidates are weakly associated with conserved DNA sequence motifs

Fungal mating depends on the transcription of mating genes controlled by master regulatory transcription factors [16, 30]. In *S. cerevisiae*, haploid cells are competent to mate because a transcription factor (TF), Ste12, binds the pheromone-responsive element (PRE motif) upstream of mating genes to induce transcription. Although we were initially attracted to the idea of using such motifs to help identify candidate pheromone genes, the specific transcription factor involved in regulating mating gene expression can change across groups of related species [31]. Specifically, Sorrells and colleagues identified the existence of Ste12 TF motifs upstream of mating genes across a large part of the budding yeasts (Saccharomycotina), however these motifs only control MAT**a**-specific gene expression in the Saccharomyces lineage.

We considered 8 pheromone-activated genes and 7 MAT**a**-specific genes essential to mating in *S. cerevisiae* expressed in haploid yeast cells that are either induced by pheromone stimulation or specifically induced in **a**-like haploid cells and found that most of these genes have homologs in the 332 sequence yeast species (Figure 4A). We then used a computational search (MEME) [32] on upstream regions (inferred promoter region) of these homologs to identify conserved TF binding sites involved in mating (Figure 4A). Very few significant motifs were identified by restricting the search to these 15 mating promoters across the full set of 332 sequenced species (Supp Figure 4).

**Figure 4.**
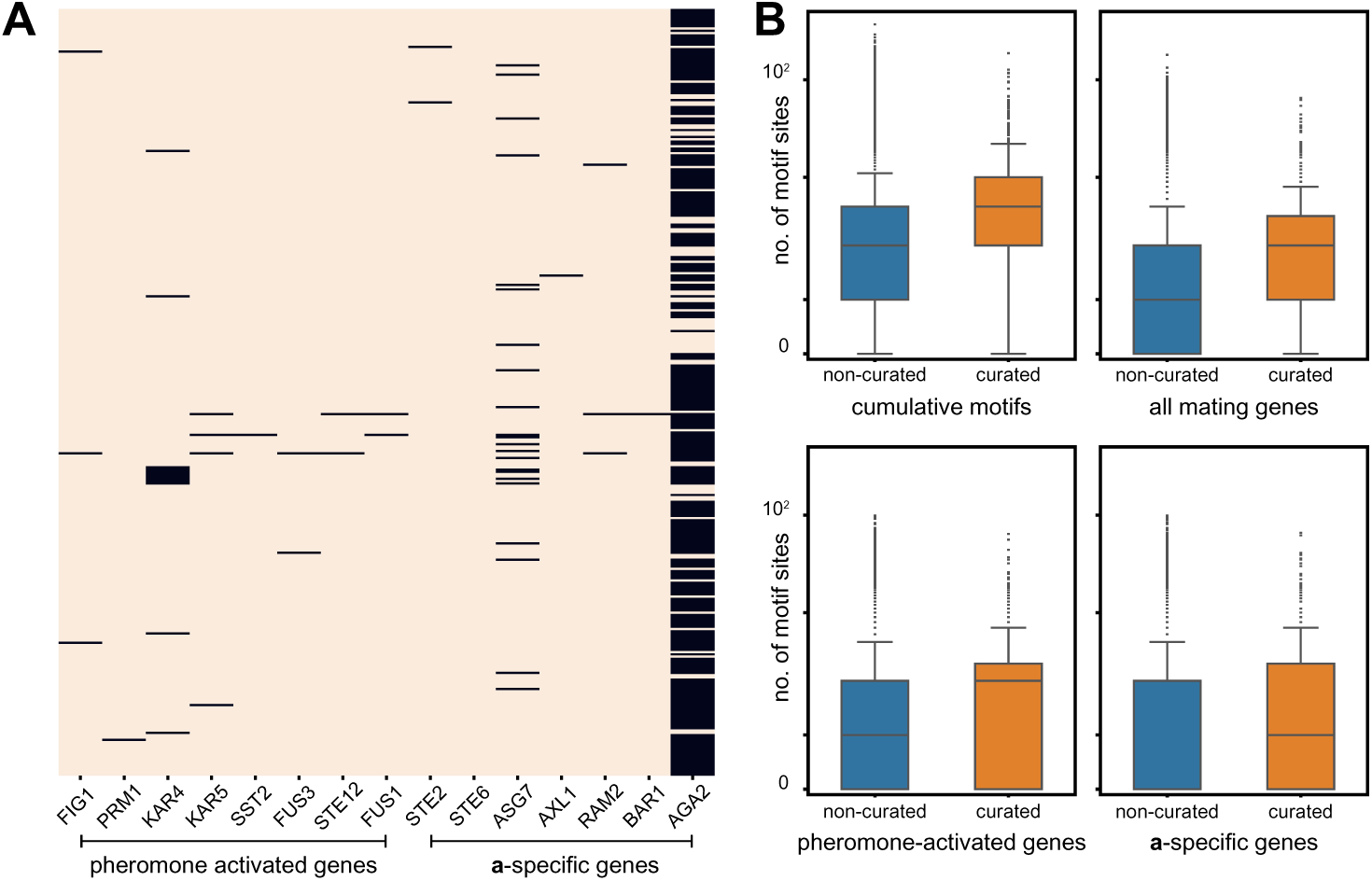
Best pheromone candidates are weakly associated with motifs that regulate other mating genes. (**A**) We considered 15 genes essential to mating in *S. cerevisiae* that are either induced by pheromone stimulation or specifically induced in **a**-like haploid cells (see labels at bottom). The figure represents the sequences of 332 yeast genomes (rows) and the detection of homologs of each of these 15 specific mating genes (columns), with a black bar showing the failure to detect a homolog of the mating gene in a particular genome. (**B**) Although true candidates are not clearly identified by the presence of mating regulatory motifs in their promoters, there is a noticeable enrichment of motifs identified using all mating genes, pheromone-activated genes, **a**-specific genes, or the sum of all identified motifs of every class (cumulative) upstream of manually curated candidates (orange) compared to all candidates that have a homologous copy in genomes of the conserved-pheromone phylogroup (blue). Each box plot represents the distribution of motif counts upstream of every candidate, with the midline representing the median, the box and whiskers representing quartiles, and the dots representing outliers. The enrichment of MEME identified motifs upstream of curated candidates are found to be significant using a two-sample Kolmogorov-Smirnov test.

Similar to pheromone homolog detection, we improved motif discovery by pooling mating genes across species within an evolutionary time horizon where the predicted regulatory motif is conserved upstream of the **a**-specific mating genes (Figure 3A) [31]. We assume that mating TF binding sites are conserved within species that are separated by a time horizon smaller than that corresponding to the divergence of *S. cerevisiae* and *V. polyspora* (Figure 3A) [31]. This yields 37 clades with at least 2 species, accounting for 290 genomes (Figure 3B). MEME identified 52, 44, and 37 significant regulatory motifs (e-value < 0.01) when all mating genes, only pheromone-activated genes, or only **a**-specific genes were pooled, respectively (Supp Figure 4). These three pooling strategies only find potential motifs in 23, 21, and 16 of the 37 phylogenetic groups where regulatory motifs are expected to be conserved, highlighting the difficulty in finding regulatory motifs using a limited set of co-regulated promoters. Twenty-five of the 37 phylogenetic groups (266 of the 290 genomes) have a predicted conserved mating TF-binding motif identified from at least one of the three sets of mating genes. Several significant motifs are variations of the binding site for the mating master regulator Ste12 of the Saccharomycetaceae clade of yeast (‘TGAAACA’). In the analyses below, we considered all identified significant MEME motifs to ensure accounting of yeast clades with divergent **a**-specific gene regulation.

We counted the presence of these identified mating TF motifs upstream of pheromone candidate genes in the presumed promoter region using the FIMO algorithm [33] and counted significant hits (p-value < 0.01). Of the 75 504 unique candidate loci across 290 species, 57 573 (76%) candidates across 266 species have at least one identified motif in their promoters. Of the 792 manually curated pheromone candidates in 238 species, 684 (86%) candidates in 216 species have at least one copy of any of the identified mating motifs in their promoter. The remaining 108 pheromone candidates are identified to have homologous pheromone candidates in related strains but do not have any identifiable regulatory motif upstream.

In summary, regulatory motifs may be useful to validate candidates identified by other approaches. But motifs are found upstream of nearly as many of the pheromone candidates prior to curation (76%) as in the manually curated set (86%) and are thus not sufficient to separate the true pheromones from the filtered [N…CAAX] candidates. Although the presence of mating-related TF-binding motifs in candidate promoters cannot be used to predict true pheromones, there is a statistical enrichment of mating-related motifs (from any or all subsets of mating genes), upstream of manually curated candidates (Figure 4B).

### Identifying the *Y. lipolytica* a-factor

The Yarrowia clade of budding yeasts is used for the heterologous expression of proteins using hydrocarbons as a metabolic carbon source [34, 35], and the production of secondary metabolites [36]. In basic research, *Y. lipolytica* is a valuable model system to study fatty acid metabolism in peroxisomes [37] and mitochondrial complex I in an obligate aerobe [38, 39]. Other than the 5 Yarrowia genomes included in the 332 genomes, we also analyzed a sixth genome available from NCBI for pheromone candidates (*Y. sp. 30695*; Figure 5A). The best candidate pheromone is present in multiple copies in each of the genomes (Figure 5A and Table 2). Comparing the flanking regions of copies in all the genomes confirms that all identified loci are unique within a genome, with pairs in the closest species pairs (*Y. divulgata* and *Y. deformans*; *Y. keelungensis* and *Y. sp. 30695*) being syntenic (Supp Figure 5C). The presumed mature peptide sequence of the pheromone is completely conserved within the clade with the exception of a single conservative mutation (F to Y; Figure 5A). Although there are several DNA polymorphisms in the mature peptide’s sequence, they are synonymous mutations (Figure 5A). In contrast, there is both non-synonymous variation and indels in the N-terminal region of the pheromone precursor in the Yarrowia lineage (Figure 5A). This is consistent with previous experiments screening for mutants in *S. cerevisiae* which showed that the sequence identity of the proteolyzed N-terminal sequence does not affect the production of mature **a**-factor [19].

**Figure 5.**
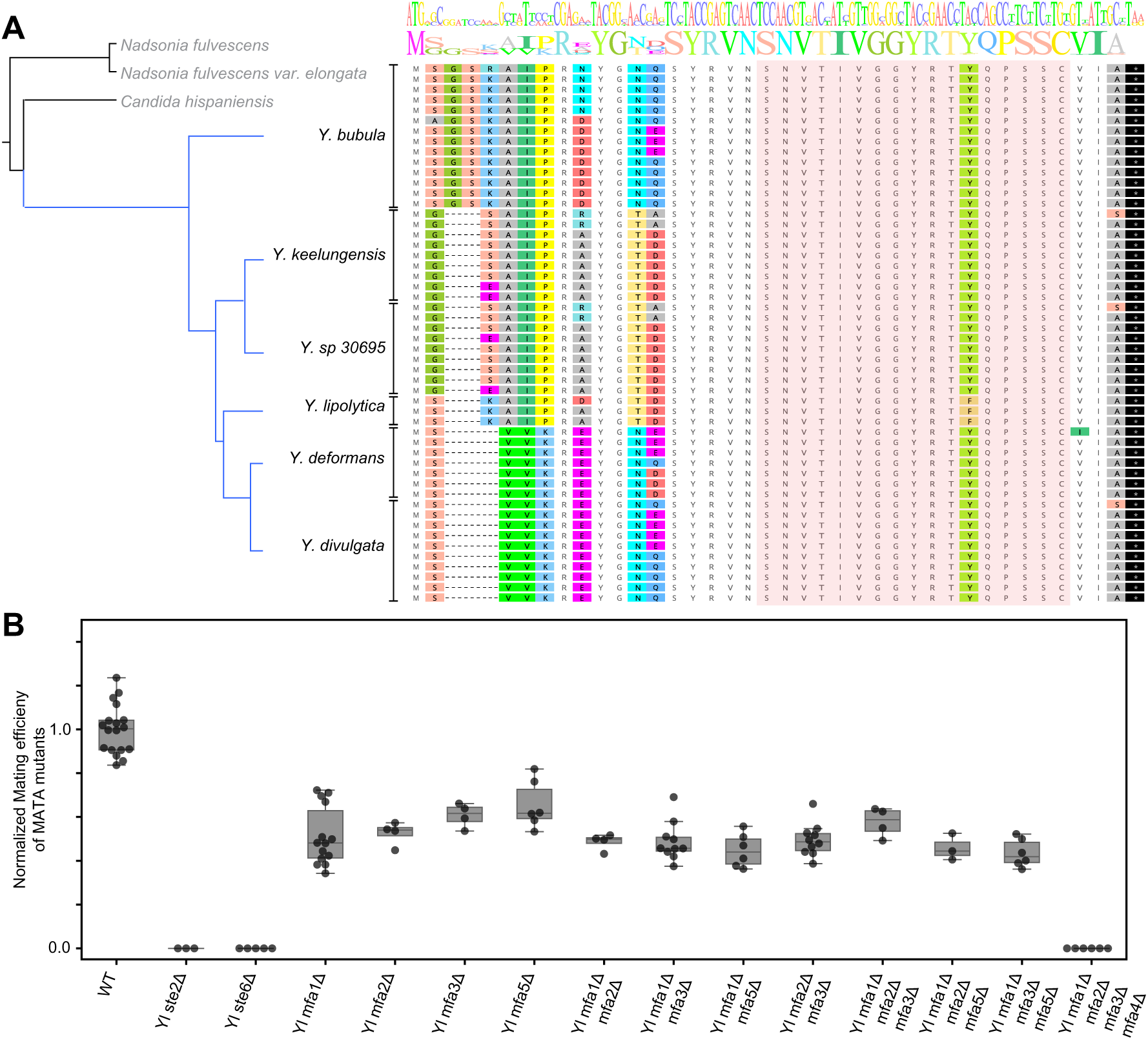
All species in Yarrowia clade of yeasts have a homologous farnesylated pheromone that is encoded in multiple copies per genome. (**A**) Phylogenetic tree of Yarrowia species (and outgroup in grey) from the set 332 of sequenced yeast of genomes [25], along with *Y. sp 30695* which is a sister species of *Y. keelungensis*. The genome of *Y. sp. 30695* was analyzed independently to identify pheromone candidates. The translated ORFs of manually curated candidates from each species are aligned and ordered according to the phylogenetic relationship between the species. The red shaded region represents the candidate mature pheromone sequence, showing no non-synonymous variation across Yarrowia, except for a conservative change of phenylalanine (F) to tyrosine (Y) in *Y. lipolytica*. At the top are alignment logos of nucleotide and amino-acid sequences, which illustrate that non-synonymous variation is enriched in the proteolyzed N-terminal fragment, compared to the mature pheromone and conserved motifs. Synonymous variation is still present in the mature pheromone sequence (see wobble bases of nucleotide logo). (**B**) The mating efficiency of MATA haploid derivatives of *Y. lipolytica* with combinations of the four pheromone loci deleted was evaluated using a semi-quantitative mating protocol. Measurement for each genotype is represented as a group of at least 3 replicate measurements performed as both biological and technical replicates. Single- double- and triple-mutants of *Yl*MFA genes show reduced mating, but only the quadruple deletion of all pheromone loci is deficient in mating to a comparable degree as the receptor (*Ylste2*Δ) and pheromone exporter (*Ylste6*Δ) deleted strains.

**Table 2.**
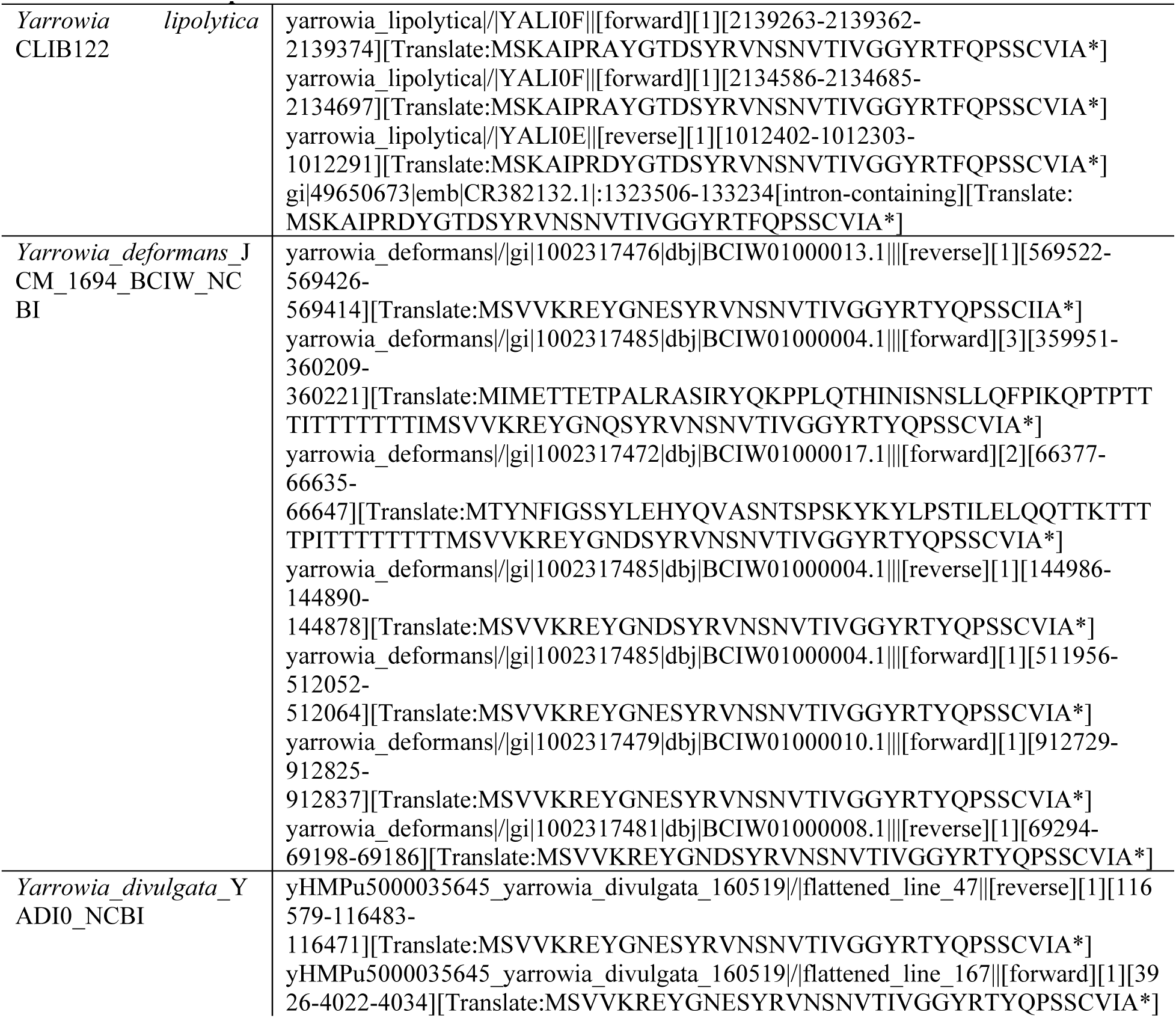
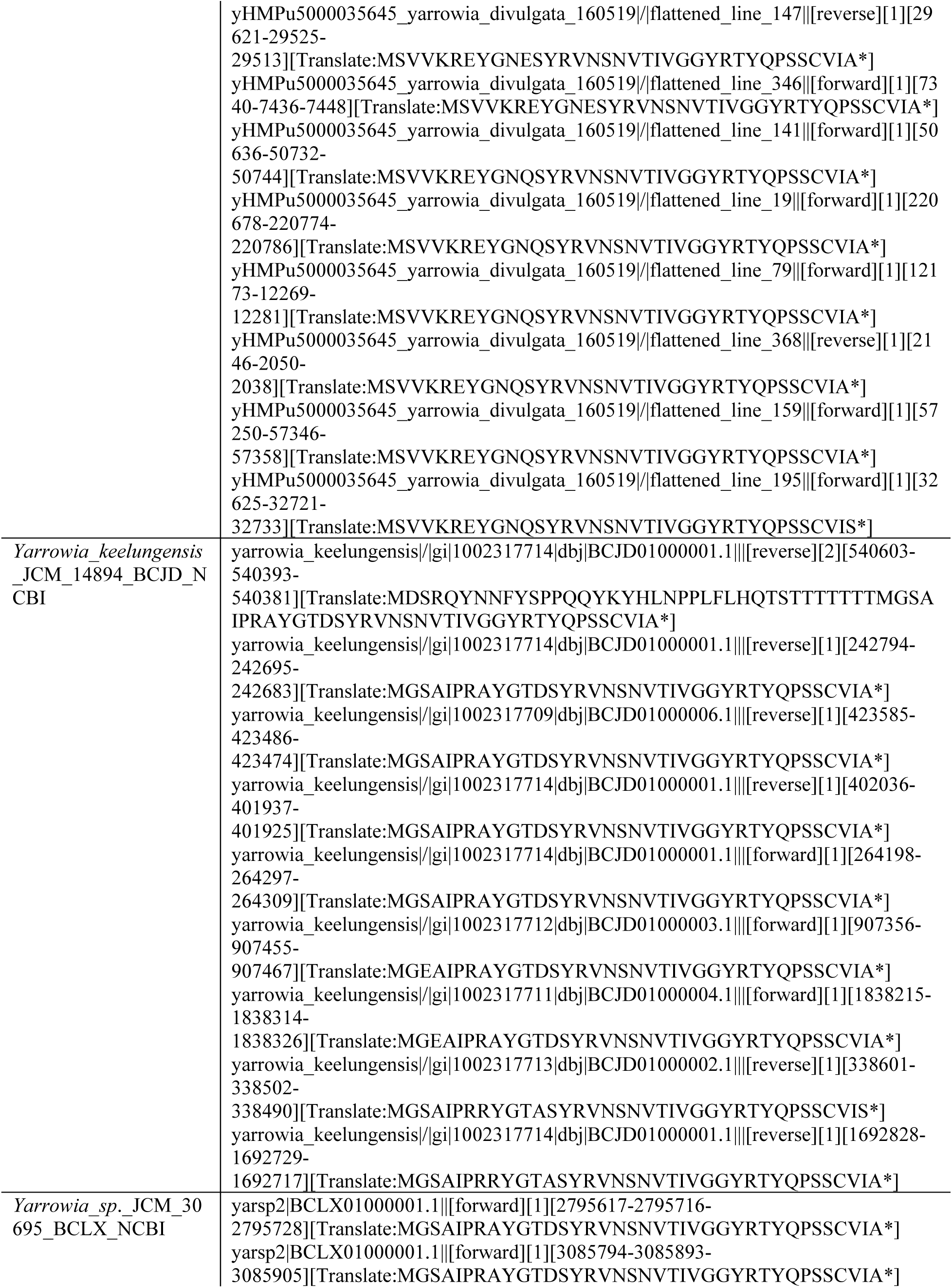
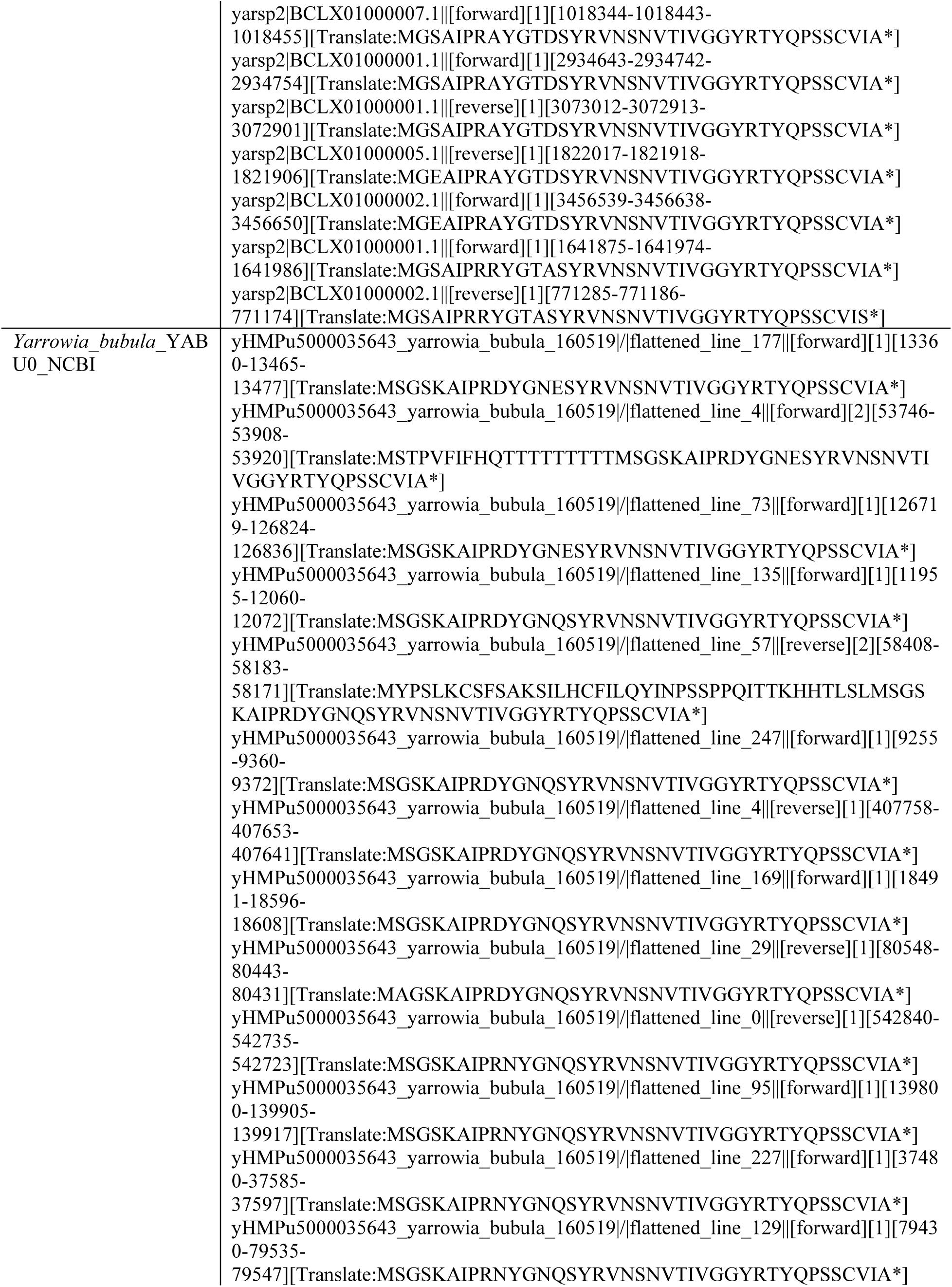
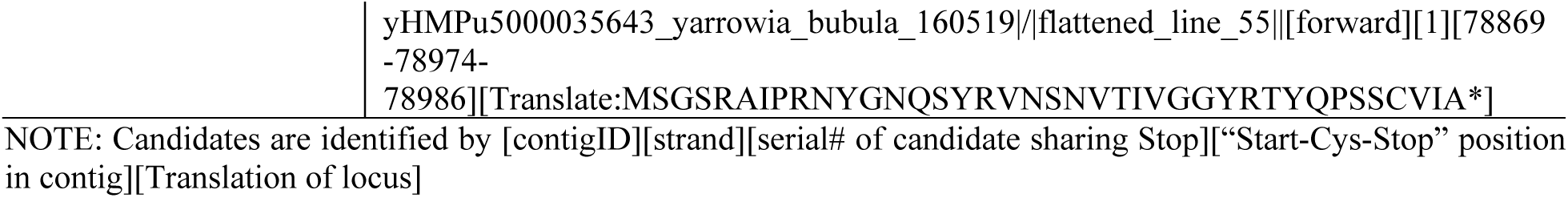
Conserved pheromone candidates in Yarrowia clade

We showed that the *Y. lipolytica* MATA mating-type is functionally homologous to MAT**a** of *S. cerevisiae*: *Ylste2Δ* (YALI0F03905g, ortholog of the α-factor receptor) and *Ylste6Δ* (YALI0E05973g, ortholog of Ste6) deletions are mating-deficient in MATA, and a *Ylste3Δ* (YALI0F11913g, ortholog of the **a**-factor receptor) deletion is mating-deficient in MATB (Supp. Figure 5A). We deleted the best pheromone gene candidates identified by the algorithm (*Yl*MFA1, *Yl*MFA2, *Yl*MFA3) in MATA as single-, double- and triple-deletions (Figure 5B). We saw a decrease in the mating efficiency in all the deletion strains, suggesting that the candidates play a role in mating. However, we were surprised to find that the triple mutant did not eliminate mating. Comparing the copy number of the pheromone candidate in different Yarrowia species revealed that *Y. lipolytica* is an outlier with the fewest pheromone copies (3) compared to other Yarrowia species (7, 9, 9, 10 and 14) (Figure 5A and Table 2). Because the triple mutant still retained mating activity, we hypothesized that there is at least one additional, intron-containing copy of the pheromone in the *Y. lipolytica* genome.

We searched translated genome fragments (using tBLASTN) for loci that might code for another copy of the best candidate protein sequence and noticed a hit on Chromosome F. This locus encodes a copy of the pheromone synonymous to another locus (*YlMFA3*) but with an intron that disrupts the ORF, masking it from our algorithm (Supp. Figure 5B). The intron matches the precise expectations of a *Y. lipolytica* intron in length, 5’ splice site (5’ ss), branch point (BP), and 3’ splice site (3’ ss) implying this is a true extra copy of the pheromone [24]. The DNA sequence of the intron-containing gene is a close homolog of one of the intron-less genes, suggesting that an intron jumped into the gene after its duplication from a genomic copy (Supp Figure 5B).

A single gene copy of the pheromone can support mating in other yeasts that mate under nutrient starvation at low efficiency like *S. pombe* [40]. We therefore made the full quadruple-deletion (*Yl*MFA1, *Yl*MFA2, *Yl*MFA3, *Yl*MFA4) and confirmed that this manipulation abrogates mating as effectively as deleting the pheromone transporter or receptor (Figure 5B). Thus, our experiments in the model yeast *Y. lipolytica* confirm that our pheromone detection algorithm can identify the true farnesylated pheromone of a yeast clade based on the conservation of maturation motifs and multiple homologous copies in closely related genomes. This algorithm will enable annotation of genomes to include the short pheromone genes across the full yeast lineage.

## Discussion

We designed an algorithm that inputs raw genome sequence and outputs candidates of farnesylated pheromones in fungi. This is of particular importance for fungi that are of medical (*C. albicans* and *C. neoformans*) [41] and agricultural (*U. maydis* (corn smut)[42], *P. striiformis* (wheat stripe rust) [43] and *M. oryzae* (rice blast) [44]) relevance, because pheromone signaling and mating have been correlated with the development of alternate cell morphologies that play a role in virulence. We experimentally validated the candidate pheromone of the Yarrowia clade by genetic deletions and mating assays in *Y. lipolytica*. Although the regulatory pathways of mating in fungi are broadly conserved [31], we found that the conservation of pheromone sequence in closely related species is a stronger predictor of pheromones than conserved regulatory sequences in the promoters of these genes. Our method is currently limited by the inability to identify candidates that contain introns, but can be expanded to include models of introns in fungi [24].

In yeasts that have been used as laboratory models (*S. cerevisiae* and *S. pombe*) or for the industrial production of proteins and secondary metabolites (*K. pastoris* and *Y. lipolytica*), pheromones can be used as tools to control the cell biology and transcriptional program as needed. In Ascomycota (specifically in yeast-like fungi), α-factor-like pheromones remain the more frequently studied pheromone signaling cascade because the genes that encode them are easily identified by the presence of multiple repeats encoded in a single prepro-peptide and the pheromone molecules are easy to produce and biochemically well-behaved. However, it is important to identify farnesylated pheromones because they represent the ancestral form of pheromones, with the Basidiomycota, such as *U. maydis* and *T. mesenterica*, only possessing farnesylated pheromones [2, 17, 45]. Our method relies not on standard homology search, but rather is an algorithmic filter that narrows down the list of all possible ORFs to a smaller list of candidates that can be tested by experimental validation. Specifically, our current version identifies most copies of a predicted pheromone gene but may not identify all copies (due to the presence of unaccounted introns, for example). This tool provides a platform to add other filters that might be relevant to specific clades of fungi, based on known pheromones of closely related fungi.

Finally, the divergence of pheromones across the yeast lineage gives us a window into the evolution of short, unstructured peptides. The peptide sequence of pheromones are presumably constrained by needing to co-evolve with their pheromone transporters [10] and receptors [9, 11]. In addition to sequence divergence over long evolutionary timescales, pheromones also show copy number variation between the genomes of sister species indicating selection for gene duplications that increase the level of pheromone secretion and thus promote efficient mating [46]. Modern methods indicate that short peptide-coding genes are common in genomes across the tree of life, but their identification and functional characterization remain difficult [47, 48]. The pheromones of the fungal lineage offer a class of short coding genes with clear physiological roles to study sequence and copy number evolution across a lineage of a billion years.

## Methods

### Strains and plasmids

*Y. lipolytica* pair of mating strains (ML16507 and ML16510) were a gracious gift from Joshua Truehart (DSM ltd) and are derivatives of the sequenced CLIB122 strain [49]. Genomic transformation was done using standard protocols [50, 51]. Gene deletions were done by replacing the ORF with an auxotrophic marker for *YlLEU2* or *YlURA3*. To re-use the URA3 marker, deletions were made with a *YlURA3* fragment flanked by repeat regions (obtained from plasmid PMB5082, a gift from Joshua Truehart) that can excise the URA3 cassette upon selection on 5FOA. Geneious Prime 2021.2.2 was used to align and compare candidate sequences.

### *Y. lipolytica* semi-quantitative mating assay

All strains listed in Table 3 were constructed from our wildtype MATA strain (ML16507) using a chemical transformation protocol reported in the literature [51]. We modified a quantitative mating protocol used in [52] to test the mating efficiency of strains with deletions of pheromone candidates against our MATB partner (ML16510). Briefly, exponential cultures in yeast extract peptone dextrose medium (YPD) of the mating gene deleted MATA strain and partner MATB strain were harvested, and 2.5 x 10^6^ cells of each partner were mixed in 150 µL sterile water + 0.02% (w/v) bovine serum albumin. Mating mixtures were transferred onto filters using a filter assembly (with the cells spreading to about 5mm radius), and the filters (with cells) were moved onto YM mating media plates (3 g/L yeast extract, 5 g/L Bacto-peptone, 5 g/L malt extract and 20 g/L Bacto-agar) [52]. These plates were incubated at 28°C in the dark for three days (70-74 h). The density of the inoculum cell suspensions was measured using a Coulter counter for normalization. After 3 days filters with the mating mixtures were moved into 3 mL YP + 2% (v/v) glycerol + 0.05% (w/v) dextrose and incubated on a roller drum at 30°C for 3 h to recover cells from filters. The cultures were transferred to microfuge tubes and sonicated to disrupt clumps, before using a Coulter counter to check the cell density to normalize for plating. The pelleted cells were resuspended in water + 0.02% (w/v) bovine serum albumin and plated on diploid selective media (CSM-Lys-Ade). The cell suspensions were counted using the Coulter counter to normalize the plating density. The mating efficiency was calculated as the number of diploid cells for a sample normalized to the number of diploid cells from the control mating (ML16507 + ML16510) performed on the same day (code on Github at https://github.com/sriramsrikant/pheromoneFinder). The experiment was repeated with biological replicates and plating replicates to account for intrinsic noise of mating mixtures.

**Table 3.** *Y. lipolytica* strains used in this study.

| *Y. lipolytica* strain label | Genotype | Source |
| --- | --- | --- |
| ML16507 | *MATA; ura3-302; leu2-270; lys1-11; Ku70Δ* | Gift from Joshua Truehart (DSM ltd.) |
| ML16510 | *MATB; ura2-6861; leu2-3; ade1; Ku70Δ* | Gift from Joshua Truehart (DSM ltd.) |
| yaliSS001 | ML16507; *Ylste2Δ:LEU2Yl* |  |
| yaliSS005 | ML16507; *Ylste6Δ:LEU2Yl* | This study |
| yaliSS007 | ML16507; *Ylmfa2Δ:URA3Yl* | This study |
| yaliSS009 | ML16507; *Ylmfa1Δ:URA3Yl* | This study |
| yaliSS011 | ML16507; *Ylmfa3Δ:URA3Yl* | This study |
| yaliSS017, yaliSS018 | ML16507; *Ylmfa1Δ:LEU2Yl, Ylmfa2Δ:URA3Yl* | This study |
| yaliSS019, yaliSS020 | ML16507; *Ylmfa2Δ:URA3Yl, Ylmfa3Δ:LEU2Yl* | This study |
| yaliSS021, yaliSS022 | ML16507; *Ylmfa1Δ:LEU2Yl, Ylmfa3Δ:URA3Yl* | This study |
| yaliSS023, yaliSS024 | ML16507; *Ylmfa2Δ:LEU2Yl, Ylmfa3Δ:URA3Yl* | This study |
| yaliSS027, yaliSS028 | ML16507; *Ylmfa1Δ* | This study |
| yaliSS029, yaliSS030 | ML16507; *Ylmfa1Δ, Ylmfa3Δ:LEU2Yl* | This study |
| yaliSS031, yaliSS032 | ML16507; *Ylmfa1Δ, Ylmfa2Δ:URA3Yl, Ylmfa3Δ:LEU2Yl* | This study |
| yaliSS035, yaliSS036 | ML16507; *Ylmfa1Δ, Ylmfa4Δ* | This study |
| yaliSS037 | ML16507; *Ylmfa1Δ, Ylmfa2Δ:LEU2Yl, Ylmfa4Δ* | This study |
| yaliSS039, yaliSS040 | ML16507; *Ylmfa1Δ, Ylmfa3Δ:LEU2Yl, Ylmfa4Δ* | This study |
| yaliSS041 | ML16507; *Ylmfa1Δ, Ylmfa2Δ:LEU2Yl, Ylmfa3Δ:URA3Yl, Ylmfa4Δ* | This study |
| yaliSS045, yaliSS046 | ML16507; *Ylmfa4Δ* | This study |
| yaliSS047 | ML16507; *Ylmfa1Δ, Ylmfa2Δ:LEU2Yl, Ylmfa3Δ, Ylmfa4Δ* | This study |

NOTE: In *Y. lipolytica*, MATA and MATB mating-types are equivalent to MATa and MATα of *S. cerevisiae* respectively.

**Table 4.** “Best” candidate pheromones identified in Y. lipolytica for experimental validation.

| Label | Expt. Label (for lab material) | Chromosomal position |
| --- | --- | --- |
| *YIMFA1* | YlPher01 | Yali0F_6\|\|[forward][2134586-2134685-2134697] |
| *YIMFA2* | YlPher02 | Yali0F_6\|\|[forward][2139263-2139362-2139374] |
| *YIMFA3* | YlPher03 | Yali0E_5\|\|[reverse][1012402-1012303-1012291] |
| *YIMFA4* | YlPher05 | Yali0F_5\|\|[forward][1323342-1323493-1323505] |

NOTE: Candidates located by [contigID][strand][serial# of candidate sharing Stop][“Start-Cys-Stop” position in contig].

### Algorithmic filter to identify pheromone candidates

As described in the main text, we brute-force identified all stop codons (TAA, TAG, TGA) in the six frames of translation of a genome that are immediately preceded by the farnesylation motif, CAAX (Supp Figure 1B). We further only considered loci that have a start codon (ATG) within 100 codons upstream of the stop codon in the same frame of translation. Presence of multiple Start (Met) codons within the region of up to 300 bases upstream of a CAAX-stop motif are accounted as multiple pheromone candidates that share a common CAAX-stop terminus and at least one internal Asn/N cleavage site (Figure 2A; 125 711 ORFs encoded in 87 326 unique CAAX-stop loci). We used the chromosomes or contigs as a natural split given that their length in fungi allows for reasonable single-core performance of the sieve for stop codons in all frames of translation. To ensure our pipeline is greedy in finding candidates we excluded concerns of chromosomal position because some fungi have poorly assembled genomes. Due to the incomplete assembly of chromosomes in many yeasts, pheromone candidates that bridge contigs are necessarily lost and cannot be reasonably identified. We distributed the calculation by contig across processors on the Odyssey cluster (Research computing, Harvard FAS).

All scripts to identify candidate loci and subsequent search for homologous pheromone candidates and mating-regulation motifs are custom-written in python. Pheromone candidate homologs are determined by measuring the pairwise sequence identity of the candidate ORFs from the potential protease site (N) to the stop codon (N(X_M_)CAAX) across species within the phylogroup where pheromones are conserved (Figure 3A). About a seventh of candidates (11 065 of 78 206 [N…CAAX] candidates in conserved-pheromone phylogroups that have at least 2 species – 300 of 332 species) have at least one other candidate that is >85% sequence identical and are considered to have a homologous copy. We further expanded the set of considered homologous pheromone candidates to about a quarter (17 125 of 78 206) by considering all [N…CAAX] candidates in conserved-pheromone phylogroups that have at least one copy >70% sequence identical. Manual curation was done by aligning these candidates grouped by phylogroup and looking for large blocks of near-identical sequences distributed across species and often present in more than one copy within a genome. Motifs involved in mating regulation were identified by using MEME [32] to look for motifs upstream of known mating genes (FIG1, PRM1, KAR4, KAR5, SST2, FUS3, STE12, FUS1 as pheromone-activated genes and STE2, STE6, ASG7, AXL1, RAM2, BAR1, AGA2 as **a**-specific genes in *S. cerevisiae*) [31] in all 332 yeast genomes. We identified at most 3 significant mating-related motifs upstream of all mating genes, pheromone-activated genes, or **a**-specific genes by grouping species into phylogroups where mating regulation is expected to be conserved (Figure 3A) [31]. FIMO was then used to identify significant hits of motifs from each phylogroup upstream of the candidates of species within that phylogroup.

Analysis and plotting of statistics of filters are written in python using Jupyter notebooks. All scripts and Jupyter notebooks are available on GitHub (https://github.com/sriramsrikant/pheromoneFinder).

## Acknowledgements

We thank the members of the Murray lab and Gaudet lab for helpful discussions. S.S. was a Howard Hughes Medical Institute International Student Research fellow. This work was funded in part by NIGMS grant R01GM120996 to R.G. and NIH grant R01GM43987 and the NSF-Simons Center for Mathematical and Statistical Analysis of Biology at Harvard (#1764269 [NSF] and #594596 [Simons]) to A.W.M.

---

**Supplementary Figure 1.**
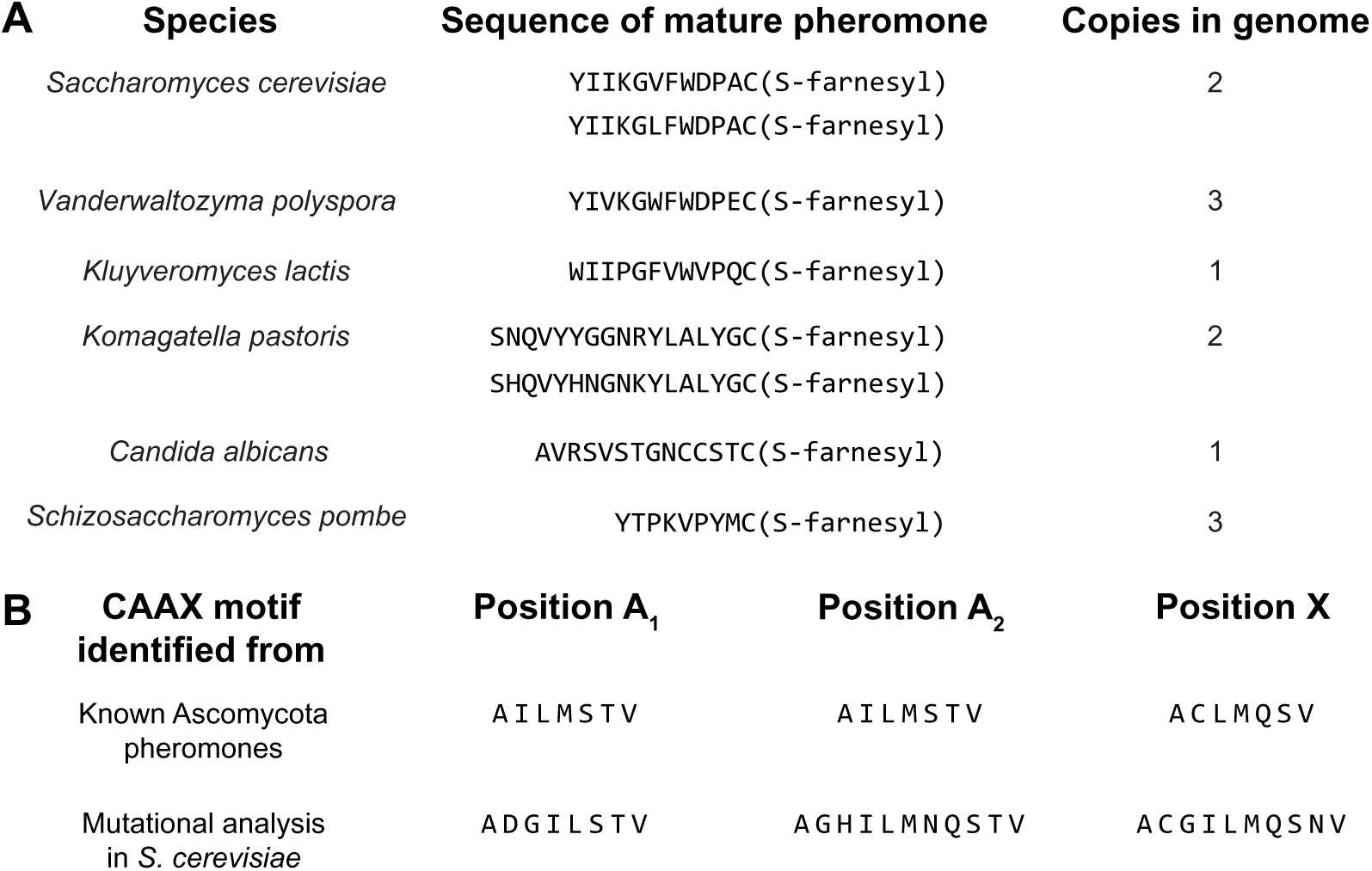
Yeast peptide pheromones share common maturation motifs. (**A**) Lipidated peptide pheromones (*Sc***a**-factor like) of yeast species (Saccharomycotina) verified by biochemistry or genetics. Yeasts have different numbers of genes in their genomes encoding identical or near-identical mature pheromones sequences. (**B**) The farnesylation [CAAX] motif is defined by cysteine that is S-farnesylated, followed by two aliphatic residues (AA) and a final variable residue (X). (Top) Known yeast pheromones provide a dictionary of possible AAX amino acid residues to identify unannotated pheromone CAAX motifs in the genome. (Bottom) To ensure that the filter is not biased by known pheromones we expanded the dictionaries used in our filter to include alternate residues that maintain farnesylation in *S. cerevisiae* [21–23]. Our final filter thus searches for all possible [CAAX-stop] motifs with A_1_, A_2_, X defined by the bottom row.

**Supplementary Figure 2.**
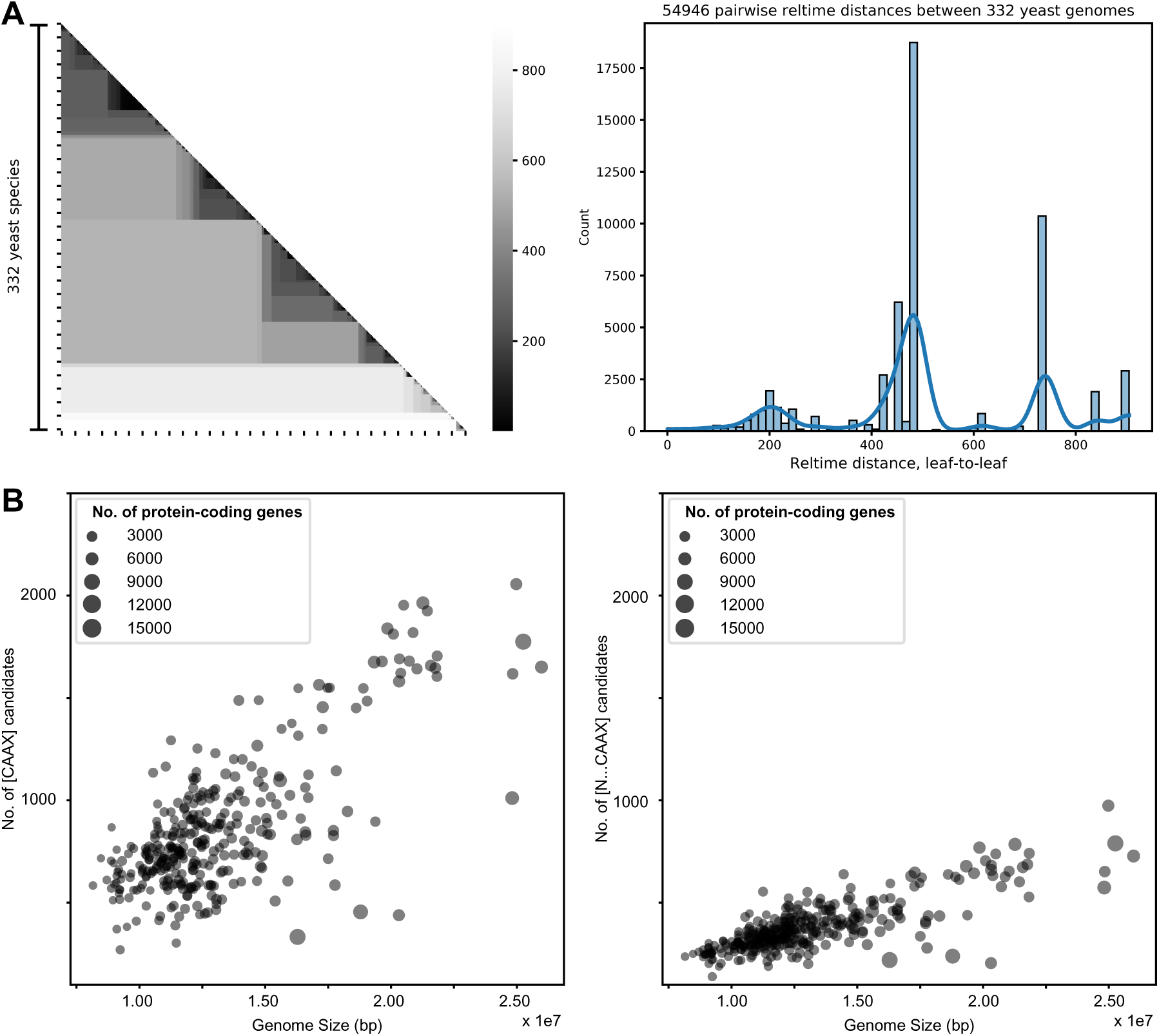
Successive filters reduce the number of candidate pheromone loci. (**A**) (Left) Matrix heatmap showing the pairwise distances amongst 332 sequenced yeast genomes evaluated using a reltime time-calibrated species divergence tree [25]. The scale bar indicates pairwise distance between extant species in millions of years. (Right) Histogram of phylogenetic distances between pairs of species from the heatmap confirms a hierarchical connection between clades, with yeasts within the same clade more related to each other than species from different clades. (B) Scatter plots of the number of [CAAX] candidates (left) and [N…CAAX] (right) candidates plotted against the genome size, where each circle represents a genome, and the diameter of the circles represents the number of annotated protein-coding genes in the genome. Successive filters in our algorithm (the presence of a C-terminal [CAAX] and an upstream, in-frame proteolytic site [N…CAAX]) reduce the number of candidate pheromone loci.

**Supplementary Figure 3.**
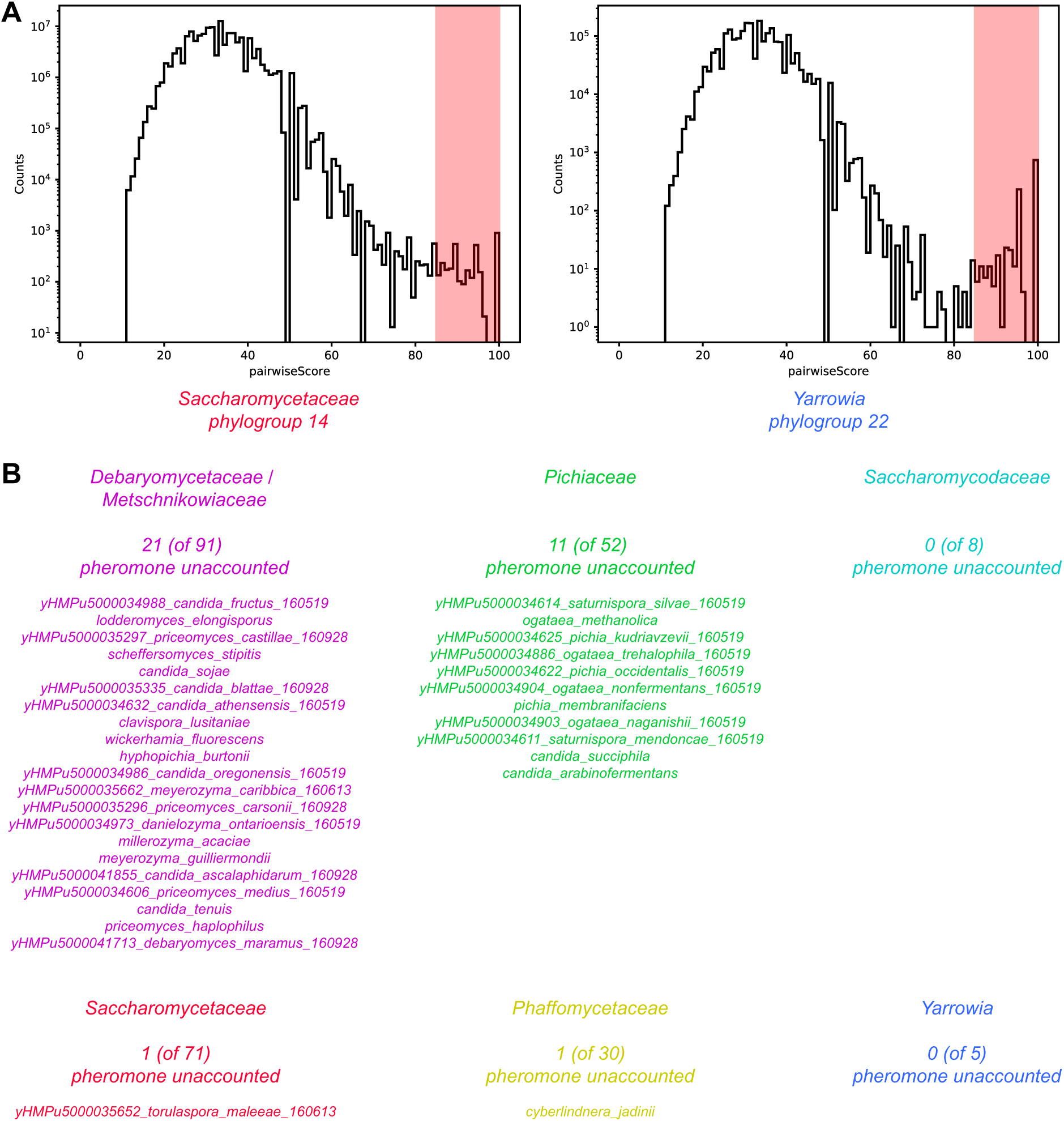
Manual curation from candidates that have homologous copies within a phylogroup identifies good pheromone candidates in most species. (**A**) Histograms of pairwise sequence identity between all [N…CAAX] candidates (calculated between potential proteolysis site to stop) identified from genomes within a phylogroup where pheromones are expected to be conserved. Because these species should have homologous pheromones, their pheromones are likely found among candidate pairs who are more than 85% identical (red shaded region). Sample histograms from phylogroup 14 (Saccharomycetaceae) and phylogroup 22 (Yarrowia) are shown. (**B**) List of species where manual curation of best pheromone candidates did not find any pheromone candidates.

**Supplementary Figure 4.**
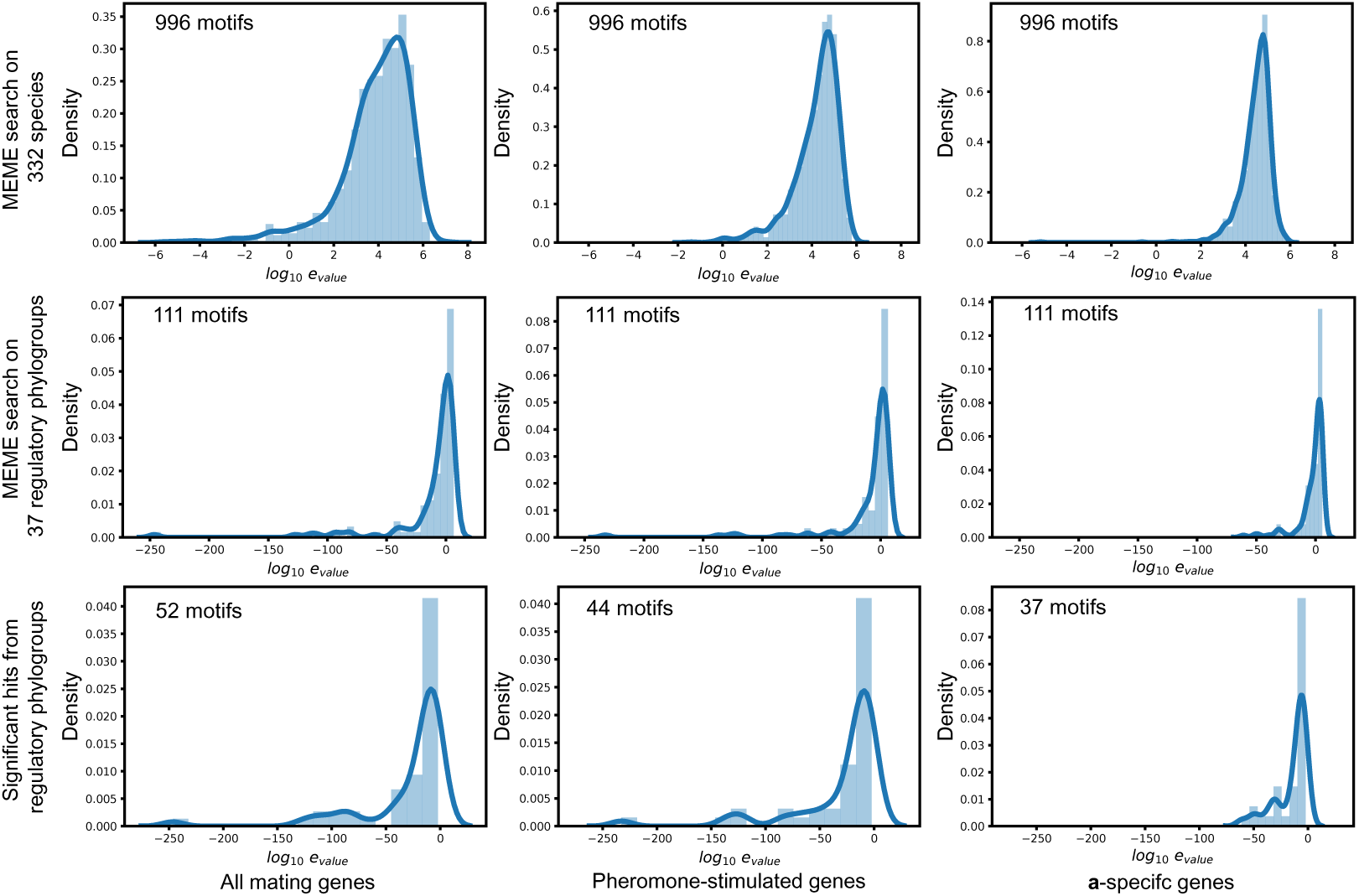
Motifs of mating regulators are found by MEME in mating genes within phylogroups. Histograms of e-values of MEME motifs identified in promoters of all mating genes (left), pheromone-activated genes (middle) or **a**-specific genes (right). (Top row) Motifs identified when analysis is done in individual species. This analysis does not identify many significant MEME motifs (e-value < 0.01). (Middle row) When mating promoters are pooled across species in conserved-mating-regulation phylogroups (290 species across 37 phylogroups), several significant motifs are identified. (Bottom row) Histograms of e-values of significant MEME motifs identified from analyzing all 290 genomes from 37 mating regulation conserved phylogroups collectively for all mating genes (52 motifs), pheromone-stimulated genes (44 motifs) and a-specific genes (37 motifs) in 266 genomes.

**Supplementary Figure 5.**
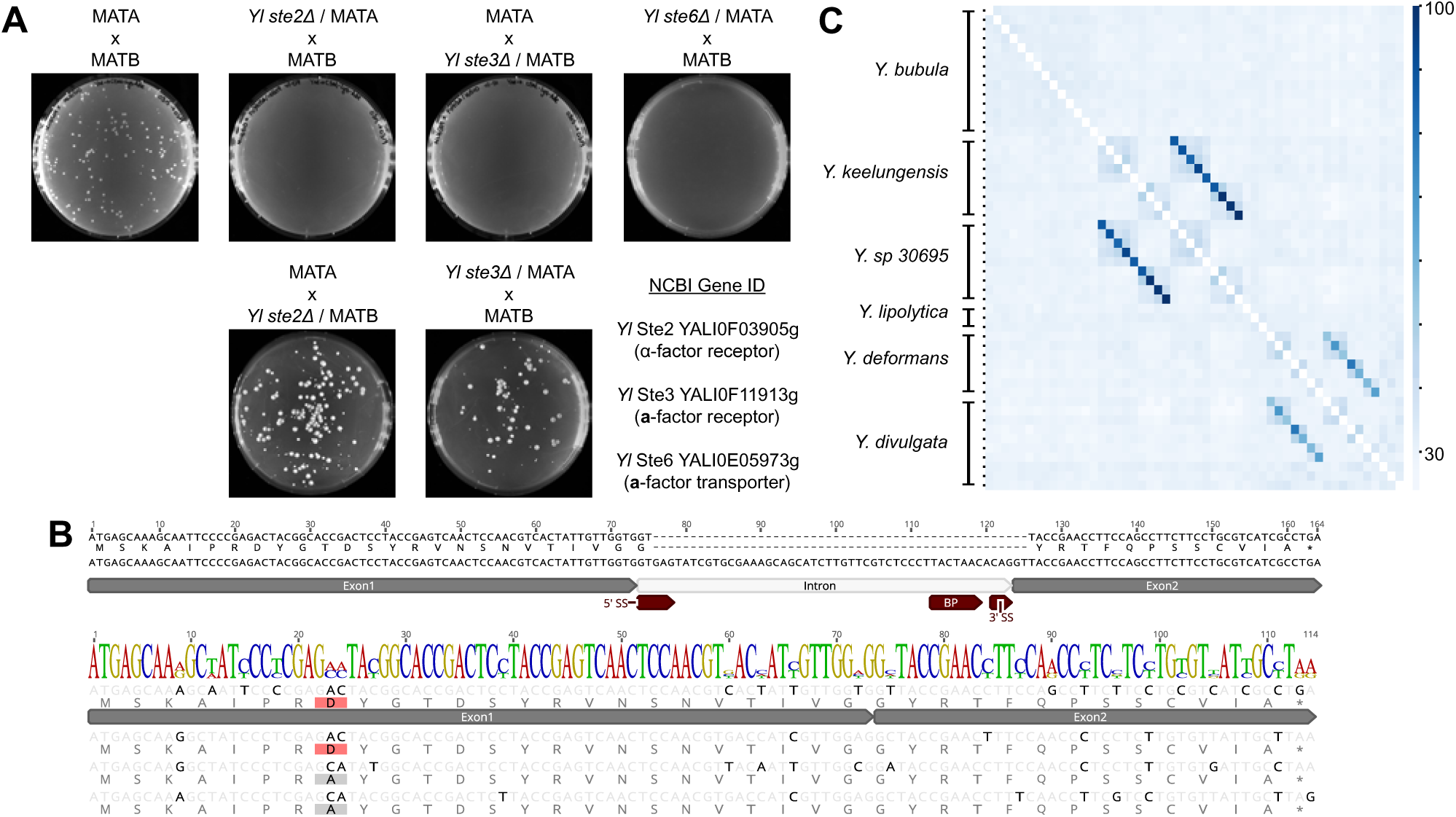
*Y. lipolytica* mating-type MATA is functionally homologous to *S. cerevisiae* MATa and MATB is functionally homologous to MATα. (**A**) Plating of *Y. lipolytica* mating mixtures from semi-quantitative mating experiments on medium on which only diploids can grow. Deletions of homologs of MAT**a** specific mating genes (α-factor receptor, Ste2 / YALI0F03905g and the **a**-factor exporter, Ste6 / YALI0E05973g) are deleterious to mating in the *Y. lipolytica* MATA haploid background. Correspondingly, deletion of a homolog of a MATα specific mating gene (**a**-factor receptor, Ste3 / YALI0F11913g) shows a mating defect in the *Y. lipolytica* MATB haploid. (**B**) *Y. lipolytica* genomic locus *YlMFA4* encodes the candidate pheromone and contains a predicted intron. (Top) Genomic locus in *Y. lipolytica* that contains a predicted intron identified by motifs (5’ ss, BP, 3’ ss) conserved across *Y. lipolytica* introns [24]. The corresponding cDNA and its translation (single-letter amino acid code) is aligned to the locus. (Bottom) The cDNA from intron-containing pheromone is aligned to the three intron-less genomic loci identified from the *Y. lipolytica* genome, showing a number of synonymous variants (darker letters) with only a single non-synonymous variant (gray and red boxes) in the N-terminal region (consistent with the pattern of variation across all Yarrowia; Figure 5A). (**C**) Aligning the 1 kbp upstream and downstream flanking regions of pheromones across the Yarrowia lineage (Figure 5A) is a measure of synteny of pheromone genes across Yarrowia genomes. Each row and column correspond to pheromone candidates in Yarrowia species with between 3 to 14 copies grouped by genome in the same order as in Figure 5A. Pairwise % identity from the alignment of the flanking regions of each locus displayed as a heat-map marking group of pheromones contained in each species. The low off-diagonal values within candidates of a species are consistent with no candidate within a species being double-counted. The high off-diagonal values all represent syntenic candidates in most-closely related pairs of species (*Y. divulgata* and *Y. deformans*; *Y. keelungensis* and *Y. sp. 30695*).

## Notes

### Competing Interest Statement

The authors have declared no competing interest.

## References

1. Lücking, R., Huhndorf, S., Pfister, D.H., Plata, E.R., and Lumbsch, H.T. (2017). Fungi evolved right on track. Mycologia 101, 810–822.

2. Jones, S.K., Jr., and Bennett, R.J. (2011). Fungal mating pheromones: choreographing the dating game. Fungal Genet Biol 48, 668–676.

3. Kurjan, J., and Herskowitz, I. (1982). Structure of a yeast pheromone gene (MF alpha): a putative alpha-factor precursor contains four tandem copies of mature alpha-factor. Cell 30, 933–943.

4. Singh, A., Chen, E.Y., Lugovoy, J.M., Chang, C.N., Hitzeman, R.A., and Seeburg, P.H. (1983). Saccharomyces cerevisiae contains two discrete genes coding for the alpha-factor pheromone. Nucleic Acids Res 11, 4049–4063.

5. Chen, P., Sapperstein, S.K., Choi, J.D., and Michaelis, S. (1997). Biogenesis of the Saccharomyces cerevisiae Mating Pheromone a-Factor. The Journal of Cell Biology 136, 251–269.

6. Michaelis, S., and Herskowitz, I. (1988). The a-factor pheromone of Saccharomyces cerevisiae is essential for mating. Molecular and Cellular Biology 8, 1309–1318.

7. Michaelis, S. (1993). STE6, the yeast a-factor transporter. Seminars in Cell Biology 4, 17–27.

8. Caldwell, G.A., Naider, F., and Becker, J.M. (1995). Fungal lipopeptide mating pheromones: a model system for the study of protein prenylation. Microbiol Rev 59, 406–422.

9. Seike, T., Nakamura, T., and Shimoda, C. (2015). Molecular coevolution of a sex pheromone and its receptor triggers reproductive isolation in Schizosaccharomyces pombe. Proceedings of the National Academy of Sciences of the United States of America 112, 4405–4410.

10. Srikant, S., Gaudet, R., and Murray, A.W. (2020). Selecting for Altered Substrate Specificity Reveals the Evolutionary Flexibility of ATP-Binding Cassette Transporters. Curr Biol 30, 1689–1702 e1686.

11. Martin, S.H., Wingfield, B.D., Wingfield, M.J., and Steenkamp, E.T. (2011). Causes and consequences of variability in peptide mating pheromones of ascomycete fungi. Mol Biol Evol 28, 1987–2003.

12. Kamiya, Y., Sakurai, A., Tamura, S., Takahashi, N., Abe, K., Tsuchiya, E., and Fukui, S. (1978). Isolation of RhodotorucineA, a Peptidyl Factor Inducing the Mating Tube Formation inRhodosporidium toruloides. Agricultural and Biological Chemistry 42, 1239–1243.

13. Kamiya, Y., Sakurai, A., Tamura, S., Takahashi, N., Abe, K., Tsuchiya, E., Fukui, S., Kitada, C., and Fujino, M. (1978). Structure of rhodotorucine A, a novel lipopeptide, inducing mating tube formation in Rhodosporidiumtoruloides. Biochemical and biophysical research communications 83, 1077–1083.

14. Sakagami, Y., Isogai, A., Suzuki, A., Tamura, S., Kitada, C., and Fujino, M. (1979). Structure of Tremerogen A–10, a Peptidal Hormone Inducing Conjugation Tube Formation inTremella mesenterica. Agricultural and Biological Chemistry 43, 2643–2645.

15. Sakagami, Y., Isogai, A., Suzuki, A., Tamura, S., Tsuchiya, E., and Fukui, S. (1978). Isolation of a Novel Sex Hormone, Tremerogen A-10, Controlling Conjugation Tube Formation inTremella mesentericaFries. Agricultural and Biological Chemistry 42, 1093–1094.

16. Bennett, R.J., and Turgeon, B.G. (2016). Fungal Sex: The Ascomycota. Microbiol Spectr 4.

17. Coelho, M.A., Bakkeren, G., Sun, S., Hood, M.E., and Giraud, T. (2017). Fungal Sex: The Basidiomycota. Microbiol Spectr 5.

18. OhEigeartaigh, S.S., Armisen, D., Byrne, K.P., and Wolfe, K.H. (2011). Systematic discovery of unannotated genes in 11 yeast species using a database of orthologous genomic segments. Bmc Genomics 12.

19. Huyer, G., Kistler, A., Nouvet, F.J., George, C.M., Boyle, M.L., and Michaelis, S. (2006). Saccharomyces cerevisiae a-factor mutants reveal residues critical for processing, activity, and export. Eukaryot Cell 5, 1560–1570.

20. Marr, R.S., Blair, L.C., and Thorner, J. (1990). Saccharomyces-Cerevisiae-Ste14 Gene Is Required for Cooh-Terminal Methylation of a-Factor Mating Pheromone. Journal of Biological Chemistry 265, 20057–20060.

21. Berger, B.M., Kim, J.H., Hildebrandt, E.R., Davis, I.C., Morgan, M.C., Hougland, J.L., and Schmidt, W.K. (2018). Protein Isoprenylation in Yeast Targets COOH-Terminal Sequences Not Adhering to the CaaX Consensus. Genetics 210, 1301–1316.

22. Stein, V., Kubala, M.H., Steen, J., Grimmond, S.M., and Alexandrov, K. (2015). Towards the systematic mapping and engineering of the protein prenylation machinery in Saccharomyces cerevisiae. PLoS One 10, e0120716.

23. Trueblood, C.E., Boyartchuk, V.L., Picologlou, E.A., Rozema, D., Poulter, C.D., and Rine, J. (2000). The CaaX Proteases, Afc1p and Rce1p, Have Overlapping but Distinct Substrate Specificities. Molecular and Cellular Biology 20, 4381–4392.

24. Neuveglise, C., Marck, C., and Gaillardin, C. (2011). The intronome of budding yeasts. C R Biol 334, 662–670.

25. Shen, X.X., Opulente, D.A., Kominek, J., Zhou, X., Steenwyk, J.L., Buh, K.V., Haase, M.A.B., Wisecaver, J.H., Wang, M., Doering, D.T., et al. (2018). Tempo and Mode of Genome Evolution in the Budding Yeast Subphylum. Cell 175, 1533–1545 e1520.

26. Adames, N., Blundell, K., Ashby, M.N., and Boone, C. (1995). Role of Yeast Insulin-Degrading Enzyme Homologs in Propheromone Processing and Bud Site Selection. Science (New York, N.Y.) 270, 464–467.

27. Dignard, D., El-Naggar, A.L., Logue, M.E., Butler, G., and Whiteway, M. (2007). Identification and characterization of MFA1, the gene encoding Candida albicans a-factor pheromone. Eukaryot Cell 6, 487–494.

28. Heistinger, L., Moser, J., Tatto, N.E., Valli, M., Gasser, B., and Mattanovich, D. (2018). Identification and characterization of the Komagataella phaffii mating pheromone genes. FEMS Yeast Res 18.

29. Seike, T., Shimoda, C., and Niki, H. (2019). Asymmetric diversification of mating pheromones in fission yeast. PLoS Biol 17, e3000101.

30. Wong Sak Hoi, J., and Dumas, B. (2010). Ste12 and Ste12-like proteins, fungal transcription factors regulating development and pathogenicity. Eukaryot Cell 9, 480–485.

31. Sorrells, T.R., Booth, L.N., Tuch, B.B., and Johnson, A.D. (2015). Intersecting transcription networks constrain gene regulatory evolution. Nature 523, 361–365.

32. Bailey, T.L., and Elkan, C. (1994). Fitting a mixture model by expectation maximization to discover motifs in biopolymers. Proc Int Conf Intell Syst Mol Biol 2, 28–36.

33. Grant, C.E., Bailey, T.L., and Noble, W.S. (2011). FIMO: scanning for occurrences of a given motif. Bioinformatics 27, 1017–1018.

34. Nicaud, J.M. (2012). Yarrowia lipolytica. Yeast 29, 409–418.

35. Liu, H.H., Ji, X.J., and Huang, H. (2015). Biotechnological applications of Yarrowia lipolytica: Past, present and future. Biotechnol Adv 33, 1522–1546.

36. Markham, K.A., and Alper, H.S. (2018). Synthetic Biology Expands the Industrial Potential of Yarrowia lipolytica. Trends Biotechnol 36, 1085–1095.

37. Dulermo, R., Gamboa-Melendez, H., Ledesma-Amaro, R., Thevenieau, F., and Nicaud, J.M. (2015). Unraveling fatty acid transport and activation mechanisms in Yarrowia lipolytica. Biochimica et biophysica acta 1851, 1202–1217.

38. Kerscher, S., Dröse, S., Zwicker, K., Zickermann, V., and Brandt, U. (2002). Yarrowia lipolytica, a yeast genetic system to study mitochondrial complex I. Biochimica et Biophysica Acta (BBA) - Bioenergetics 1555, 83–91.

39. Parey, K., Brandt, U., Xie, H., Mills, D.J., Siegmund, K., Vonck, J., Kuhlbrandt, W., and Zickermann, V. (2018). Cryo-EM structure of respiratory complex I at work. eLife 7.

40. Kjaerulff, S., Davey, J., and Nielsen, O. (1994). Analysis of the structural genes encoding M-factor in the fission yeast Schizosaccharomyces pombe: identification of a third gene, mfm3. Mol Cell Biol 14, 3895–3905.

41. Boral, H., Metin, B., Dogen, A., Seyedmousavi, S., and Ilkit, M. (2018). Overview of selected virulence attributes in Aspergillus fumigatus, Candida albicans, Cryptococcus neoformans, Trichophyton rubrum, and Exophiala dermatitidis. Fungal Genet Biol 111, 92–107.

42. Chacko, N., and Gold, S. (2012). Deletion of the Ustilago maydis ortholog of the Aspergillus sporulation regulator medA affects mating and virulence through pheromone response. Fungal Genet Biol 49, 426–432.

43. Zhu, X., Liu, W., Chu, X., Sun, Q., Tan, C., Yang, Q., Jiao, M., Guo, J., and Kang, Z. (2018). The transcription factor PstSTE12 is required for virulence of Puccinia striiformis f. sp. tritici. Mol Plant Pathol 19, 961–974.

44. Li, Y., Que, Y., Liu, Y., Yue, X., Meng, X., Zhang, Z., and Wang, Z. (2015). The putative Ggamma subunit gene MGG1 is required for conidiation, appressorium formation, mating and pathogenicity in Magnaporthe oryzae. Curr Genet 61, 641–651.

45. Raudaskoski, M., and Kothe, E. (2010). Basidiomycete mating type genes and pheromone signaling. Eukaryot Cell 9, 847–859.

46. Moore, T.I., Chou, C.S., Nie, Q., Jeon, N.L., and Yi, T.M. (2008). Robust spatial sensing of mating pheromone gradients by yeast cells. PLoS One 3, e3865.

47. Couso, J.P., and Patraquim, P. (2017). Classification and function of small open reading frames. Nat Rev Mol Cell Biol 18, 575–589.

48. Andrews, S.J., and Rothnagel, J.A. (2014). Emerging evidence for functional peptides encoded by short open reading frames. Nat Rev Genet 15, 193–204.

49. Dujon, B., Sherman, D., Fischer, G., Durrens, P., Casaregola, S., Lafontaine, I., De Montigny, J., Marck, C., Neuveglise, C., Talla, E., et al. (2004). Genome evolution in yeasts. Nature 430, 35–44.

50. Burke, D., Dawson, D., and Stearns, T. (2000). Methods in Yeast genetics: A Cold Spring Harbor Laboratory course manual, (Cold Spring Harbor Laboratory Press).

51. Verbeke, J., Beopoulos, A., and Nicaud, J.M. (2013). Efficient homologous recombination with short length flanking fragments in Ku70 deficient Yarrowia lipolytica strains. Biotechnol Lett 35, 571–576.

52. Rosas-Quijano, R., Gaillardin, C., and Ruiz-Herrera, J. (2008). Functional analysis of the MATB mating-type idiomorph of the dimorphic fungus Yarrowia lipolytica. Curr Microbiol 57, 115–120.

